# Translesion synthesis polymerase κ is recruited to ribosomal DNA to maintain rRNA transcription

**DOI:** 10.1101/2022.10.27.513845

**Authors:** Shilpi Paul, Abbey Rebok, Paolo Cifani, Anirban Paul, Darryl Pappin, Tony T. Huang, Thomas E. Spratt

**Affiliations:** Department of Neuroscience and Experimental Therapeutics, Penn State College of Medicine, Hershey, PA, 17033, USA; Department of Molecular and Precision Medicine, Penn State College of Medicine, Hershey, PA, 17033, USA; Cold Spring Harbor Laboratory, Cold Spring Harbor, NY, 11724, USA; Biochemistry and Molecular Pharmacology, New York University School of Medicine, NY, 10016 USA

## Abstract

DNA polymerase κ (Polκ) is a Y-family translesion synthesis (TLS) polymerase that bypasses bulky DNA lesions, such as those induced by the carcinogen benzo[*a*]pyrene diol epoxide (BPDE). Although TLS polymerases are increasingly recognized to function at the intersection of replication, repair, and transcription, mapping their activity to specific genomic loci has remained technically challenging because all DNA polymerases incorporate canonical nucleotides. To overcome this obstacle, we employed the selective Polκ substrate *N*^2^-(4-ethynylbenzyl)-2′-deoxyguanosine (EBndG) to capture proteins associated with Polκ - synthesized DNA. Polκ was unexpectedly found enriched in the nucleolus, and Polκ - synthesized DNA is enriched with nucleolar proteins. DNA damage induced by BPDE triggered hallmarks of nucleolar stress, stabilized Polκ, and increased its retention and activity on nucleolar DNA, including ribosomal DNA (rDNA). Additional nucleolar stressors, including mitomycin C, cisplatin, and actinomycin D, elicited similar effects. The Polκ activity was independent of cell-cycle phase but required PCNA ubiquitination and was regulated by

PARP1. These findings uncover novel interactions of Polκ with nucleolar proteins, demonstrate its direct activity on rDNA, and establish its role in sustaining rRNA synthesis after DNA damage, thereby protecting cells from nucleolar stress and linking TLS polymerase function to nucleolar homeostasis.

**KEY POINTS:**

- Polκ is the first TLS polymerase shown to be active in the nucleolus and on ribosomal DNA.
- Carcinogen BPDE induces nucleolar stress and recruits Polκ to rDNA to sustain rRNA synthesis.
- Polκ nucleolar activity requires PCNA ubiquitination and PARP1, linking TLS to rDNA homeostasis.

**GRAPHICAL ABSTRACT:** 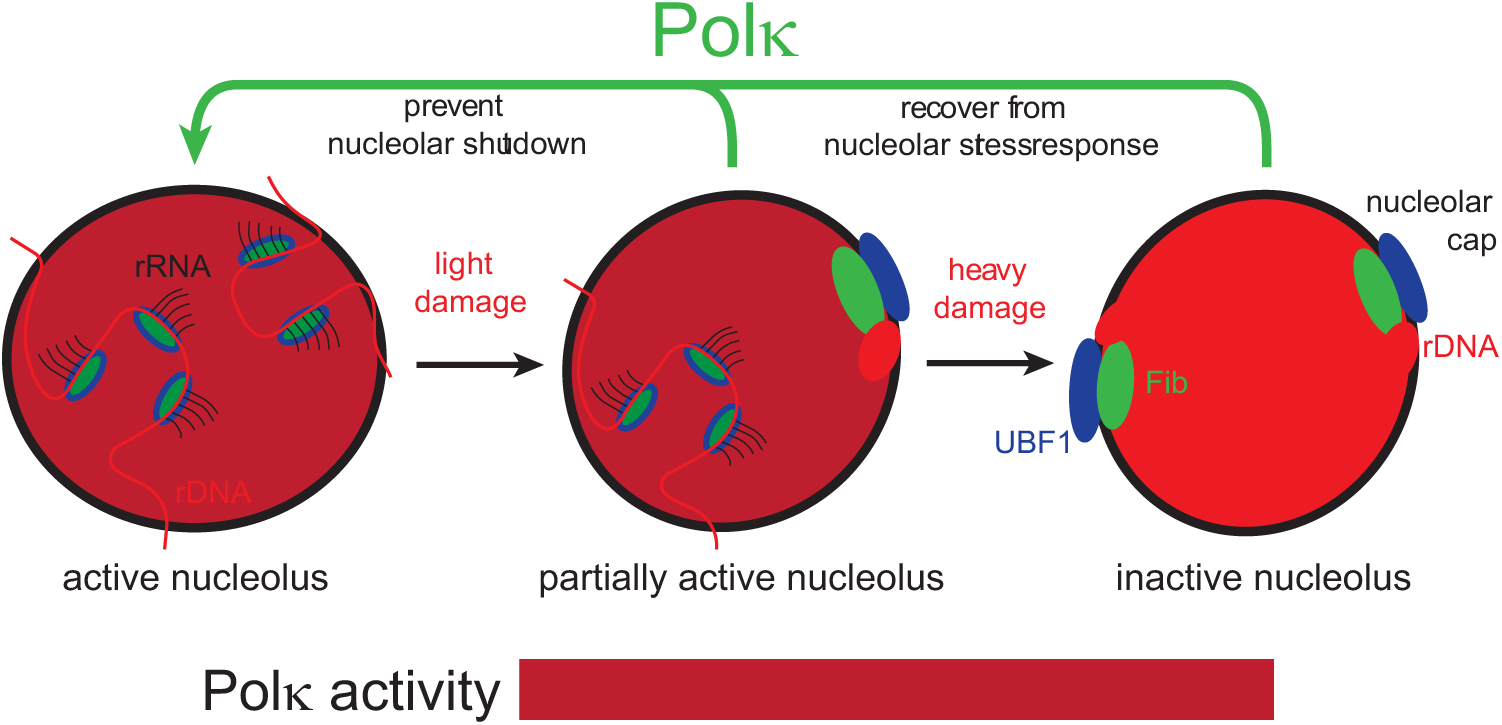

## INTRODUCTION

DNA polymerase κ (Polκ) is one of the sixteen DNA-dependent DNA polymerases expressed in humans ^1^. The most-studied function of Polκ is translesion DNA synthesis (TLS), which is one of the mechanisms by which the cell responds to DNA damage that cannot be replicated by high fidelity DNA polymerases. The carcinogen (+/-)-*trans*-benzo[*a*]pyrene-7,8-dihydrodiol-9,10-epoxide (BPDE) covalently reacts with deoxyguanosine generating 10-(deoxyguanosin-*N*^2^-yl)-7,8,9-trihydroxy-7,8,9,10-tetrahydrobenzo[*a*]pyrene (*N*^2^-BP-dG) ^2^, which Polκ bypasses efficiently and accurately by inserting dCMP opposite the lesion ^3–7^. Polκ is recruited to stalled replication forks by RAD18-mediated PCNA monoubiquitination and is essential for recovery from BPDE-induced S-phase checkpoint arrest ^8^. While the TLS function of Polκ has been the most-studied, other functions of Polκ are being elucidated. Polκ has been linked with nucleotide excision repair ^9,10^, cross-link repair ^11,12,13^, the production of short oligodeoxynucleotides in the ATR activation pathway ^14^, in replication of repetitive DNA sequences ^15,16^, and in the replication of open DNA regions early in S-phase ^17,18^. A hurdle with determining when and where a specific DNA polymerase is active is that all polymerases utilize undamaged DNA and the four deoxynucleotide triphosphates as substrates. To overcome this impediment, we utilized *N*^2^-4-ethynylbenzyl-2ʹ-deoxyguanosine (EBndG), a non-natural nucleoside that has high specificity for Polκ over other mammalian DNA polymerases_19_. Using this tool, we detected Polκ activity within nucleoli the most transcriptionally demanding compartment of the human genome.

The nucleolus forms around nucleolus organizer regions (NORs), whose major components are rDNA arrays located on the short p-arms of the five human acrocentric chromosomes 13, 14, 15, 21, and 22 ^20^. Interestingly, a subset of acrocentric p-arms lack rDNA, yet their NORs still associate with nucleoli, suggesting rDNA-independent mechanisms of nucleolar association ^21^. Human rDNA exists in hundreds of copies, with the 5.8S, 18S, and 28S rDNA sequences encoded within these NOR arrays. Transcriptionally active rDNA is organized as open chromatin and transcribed by RNA polymerase I (RNAPI) to generate a 47S pre-rRNA precursor, which is processed and assembled into ribosomes in the cytoplasm. rDNA represents the most transcriptionally active portion of the genome, and ribosome biogenesis accounts for the majority of a cell’s energy expenditure, as each rRNA molecule is used only once to produce 2,000–10,000 ribosomes per minute ^22–24^. This extraordinarily high demand for rRNA must be met continuously to sustain cell growth and proliferation ^25,26^.

The nucleoli are dispersed during mitosis, and then small nucleoli containing individual NORs are reformed during telophase when rRNA transcription by RNAPI resumes. With progression of the cell cycle, nucleoli fuse and form larger nucleoli containing multiple NORs. The nucleolus is a "tripartite" nuclear structure with multiple sub-compartments, the fibrillar center (FC), the dense fibrillar component (DFC) and the granular component (GC) ^27,28^. Transcription of pre-ribosomal RNA (pre-rRNA) occurs at the interface of the FC and DFC compartments.

The FC is enriched in components of the RNAPI machinery, such as upstream binding transcription factor (UBF1), DFC contains pre-rRNA processing factors such as fibrillarin (FBL), and the GC, where preribosome subunit assembly occurs, contains nucleophosmin (NPM1).

The intense transcriptional activity of rDNA creates unique vulnerabilities. Conflicts between RNAPI transcription and DNA replication forks generate replication stress and R-loops — stable RNA:DNA hybrids formed when nascent RNA re-anneals to the template strand, displacing the non-template DNA as single-stranded DNA (ssDNA)^29–31^. R-loops are particularly abundant at rDNA owing to its high GC content, open chromatin, and extreme transcriptional activity. Persistent R-loops and replication stress cause fork stalling and collapse, generating single-strand breaks (SSBs) that can convert to double-strand breaks (DSBs), making rDNA genes acutely vulnerable to genotoxic damage ^32^. Critically, the displaced ssDNA within R-loop structures is exposed and unprotected, rendering it susceptible to chemical carcinogens such as BPDE. As we report here, RNase H treatment — which degrades the RNA strand of RNA:DNA hybrids and thereby resolves R-loops — abolishes Polκ-synthesized DNA signal in the nucleolus, implicating R-loops as sites of Polκ activity at rDNA.

DNA damage in the nucleolus activates a coordinated nucleolar stress response. ATM/ATR activation inhibits RNAPI transcription, triggering dramatic reorganization of nucleolar structure, protein composition, and function ^33^. A hallmark of this response is the formation of nucleolar caps, in which the FC and DFC — together with rDNA — segregate to the nucleolar periphery, where DNA repair proteins concentrate ^34–36^. NPM1, a chaperone protein that is involved in ribosome biogenesis is also released into the nucleoplasm upon stress where it binds to MDM2 and prevents the MDM2-initiated degradation of p53, leading to the activation of the p53 DDR pathway ^37,38^. PARP1, nearly half of which resides in the nucleolus under basal conditions, is similarly released to sensitize cells to damage-induced apoptosis ^39,40^.

Multiple repair pathways — HR, NHEJ, and GG-NER (but not TC-NER) — operate on rDNA _34,36,41,42_ and over 4,500 proteins have been identified in the nucleolus, including 166 DNA damage response proteins ^43–46^. The nature of the DNA damage that causes the nucleolar stress response is not fully known, as DSBs, acrolein and oxaliplatin induce nucleolar cap formation while cisplatin does not ^11,47–49^. Whether TLS polymerases participate in this nucleolar damage response has not previously been investigated.

In this study we combined EBndG-based activity profiling with proteomics (iPolKD-MS), super-resolution confocal microscopy, rDNA FISH, and cell-based assays to investigate the function of Polκ in the nucleolar stress response. We identified that similar to the treatment with DNA damaging agent mitomycin C or RNAPI inhibition by actinomycin D, the carcinogen BPDE causes cells to undergo nucleolar stress. During the stress nucleolar caps are gradually formed, NPM1 is translocated from nucleolus to the nucleoplasm and the transcription of 47S pre-RNA is inhibited. Despite reduced POLK mRNA levels after BPDE-induced nucleolar stress, levels of Polκ remain high and enhanced Polκ activity is observed in the nucleolus and on rDNA. Mechanistically, Polκ nucleolar activity requires PCNA ubiquitination and its ubiquitin-binding zinc finger (UBZ) domain, is regulated by PARP1-mediated PARylation, and is independent of cell-cycle phase. Depletion of Polκ impairs recovery of rRNA synthesis following genotoxic stress. Together, these findings establish Polκ as a novel participant in the nucleolar DNA damage response and link TLS polymerase function to the maintenance of nucleolar homeostasis.

## MATERIALS AND METHODS

### Cell culture, drug treatments and plasmid transfection

HEK293, 293T-K164R ^50^, U2OS and U2OS sgPolK ^51^ cell lines were cultured in Dulbecco’s modified Eagle medium (DMEM), supplemented with 10% fetal calf serum (FCS), GM12878 cells were cultured in Roswell Park memorial Institute (RPMI) 1640 medium with 15% FCS. Both the media contained 2 mM L-glutamine and penicillin–streptomycin antibiotics and cells were grown under standard cell culture conditions in a CO_2_ incubator (37 °C; 5% CO_2_). BPDE (obtained from the Organic Synthesis Core of the Penn State Cancer Institute) was diluted in DMSO concentrations to prepare 0.1, 0.5 or 2 μM from stock by diluting and added immediately to the media. The cells with BPDE added media were incubated for 2 h in a CO_2_ incubator. For olaparib treatment, cells were incubated for 72 h with 4 μM olaparib (Sigma-Aldrich, gift from George Lucian Moldovan’s lab) added to DMEM. BPDE damage for 2 h was performed in 72 h of olaparib treated cells. For MMC and cisplatin treatment, U2OS cells were incubated with either 30 μM mitomycin C (dissolved in DMSO), 10 μM cisplatin (dissolved in 0.9% NaCl), or the solvent controls for 2 h. For actinomycin D treatment cells were incubated with 5 nM actinomycin D or DMSO for 6 h.

Transient plasmid WT-PolK GFP ^51^ transfections were performed using FuGene six reagent (Promega), according to manufacturer’s instructions. Analyses were done between 24–48 h after plasmid transfection.

### Isolation of protein on nascent Polκ synthesized DNA (iPolKD)

This procedure is a modification of the native iPOND procedure ^52^. GM12878 cells (∼8X10^6^) after treatment with 0, 0.5 μM or 2 μM for 2 h were incubated with 20 μM EdU (15 min) or 100 μM EBndG (30 min) and harvested by centrifugation at 200xg for 7.5 mins. HEK293 cells (∼4-6x10^8^) were incubated after BPDE damage with EdU (15 min) or EBndG (180 min). U2OS cells were incubated after 2 μM BPDE damage 100 μM EBndG for 30 mins and immediately harvested (sample "Pulse"). From another set the media containing EBndG was removed and fresh media added and incubated for 180 min then harvested (sample "Wash-Chase"). Sample for "Beads control" had no nucleotides (EdU or EBndG) added to the cells. Pellets were resuspended in 10 mL of pre-chilled freshly prepared NEB (20 mM HEPES-KOH pH 7.2, 50mM NaCl, 3 mM MgCl2, 300 mM sucrose, 0.5% NP 40 substitute or TX100) and rotated at 4°C for 15 mins. The nuclei were pelleted by centrifugation for 10 min at 500xg at 4°C to pellet. The nuclei pellets were resuspended in 10 mL ice-cold PBS with a serological pipette to wash (10 μL used to check the nuclei under microscope) and collected by centrifugation at 500×g for 10 min at 4°C. The nuclei pellets were completely resuspended in 10 mL of freshly prepared click reaction mixture [1 mM CuSO_4_, 2 mM tris(3-hydroxypropyltriazolylmethyl)amine (THPTA), 10 μM azido-PEG3-biotin (Alfa Aesar #J 64996) 10 mM sodium ascorbate, complemented with PBS] and incubated for 1.5 h at room temperature (RT). The nuclei were centrifuged for 10 min at 500xg at 4°C and the pellet washed 3X with 10 mL ice-cold PBS. The nuclei pellet was either immediately frozen in liquid nitrogen and kept in -80°C or were immediately lysed as described below.

The nuclei pellets were resuspended in 0.5 mL of ice-cold freshly prepared buffer B1 (25 mM NaCl, 2 mM EDTA, 50 mM Tris-HCl-pH8, 1% NP-40 substitute or TX100) supplemented with protease inhibitors (Roche cOmplete Protease Inhibitor Cocktail) and transferred to a 1.5 mL tube and rotated at 4°C for 30min. The nuclei were pelleted by centrifuged at 500xg for 10 mins at 4°C supernatant saved as low soluble chromatin and nucleoplasm fraction. The pellets were resuspended again in 0.5 mL of ice-cold buffer B1 and rotated at 4°C for 30 min. The chromatin was solubilized with a microtip sonicator (Fisher Scientific Model 500 Sonic Dismembrator) by keeping the tubes on ice and performing 6 rounds of 2 X10s on, 10s off (∼20A with 1min Rest output). The samples were centrifuged for 10 min at 16100xg at 4°C and the solubilized chromatin fraction was collected. An aliquot was saved as an input sample. The rest of the sample was supplemented to 1 mL with ice-cold buffer B1 containing protease inhibitors, 50 μL of MyOne Streptavidin T1 Dynabeads (MyOne Streptavidin T1 65602, Thermo-Fisher) were added, and rotated overnight at 4°C. The beads were washed 6 times with buffer B1. The protein was eluted by boiling in 1X Laemmli buffer.

### Mass Spectrometry and Proteomics data analysis

The eluted proteins were loaded on a 10% SDS-PAGE gel, resolved for approximately 5mm, and Coomassie stained. Gel bands containing the protein samples were excised from the gel, reduced using 3 mM tris(2-carboxyethyl)phosphine (TCEP) in 100 mM ammonium bicarbonate, and cysteine residues were alkylated using 3 mM 2-carboxyethyl methanethiosulfonate (CEMTS). Proteins were then in-gel proteolyzed using Sequencing grade modified porcine trypsin (Promega), and peptides were eluted (80% acetonitrile, 1% TFA) and lyophilized. Peptides were resuspended in 30 μL 0.1% aqueous formate, and 2 μL of solution were used for LC/MS analysis. Using an easyLC1200 nano-chromatograph (Thermo Scientific), peptides were loaded on a 20 cm x 75 μm ID column packed in house with Reprosil C18 silica particles (1.9 μm diameter, Dr. Maischt), and resolved on a 5-35% acetonitrile gradient in water (0.1% formate) at 250 nl/min.

Eluting peptides were ionized by electrospray (2000 V) and transferred into a Thermo Lumos Orbitrap mass spectrometer whose ion transfer tube was maintained at 305 °C. The MS was set to collect 120,000 resolution orbitrap precursor scans (profile mode) every 3 s (m/z 380-2000 Da, AGC= 400,000 ions, maximum injection time = 50 ms, RF lens = 30%). Precursor ions were selected for data-dependent acquisition in cycle-time mode (i.e. performing as many MS2 as possible within the 3 s interval between consecutive MS1 scans), applying the following filters: dynamic exclusion 30 s with 10 ppm mass tolerance, monoisotopic ion selection based on isotopic pattern expected for peptides, charge state 2-7 (ions with undetermined charge state were excluded), and minimum ion intensity 5,000. Precursor ions were selected using the quadrupole (isolation window = 1.2 Da), accumulated for up to 50 ms (AGC= 30,000 ions), and fragmented by HCD at stepped 26,32,38% normalized energy.

Spectra were recorded in centroid mode using the linear trap at “normal” (unit) resolution, with first mass locked to 100 Da. The reversed phase column was washed after each sample injection to prevent carry over.

Files searched using the Mascot scoring function within ProteomeDiscoverer v.2.4, with mass tolerance set at 5 ppm for MS1, and 0.5 Da for MS2. Spectra were matched against the UniProt human database (as of January 2021, containing 40,967,805 residues and 101,014 protein sequences) and a database of common contaminants (cRAP). M-oxidation, and N/Q-deamidation, were set as variable modifications. CEMTS adduct on cysteine was set as fixed modification. Trypsin was set as protease with up to 3 missed cleavages allowed. Peptide-spectral matches were filtered to maintain FDR<1% using Percolator. Extracted ion chromatograms of precursor ions intensities were integrated for label-free quantification (LFQ_ across samples and used as quantitative metric for protein relative quantification.

Proteins with peptide count 2 and above were further analyzed. The ribosomal proteins (RPL proteins) were removed from all data sets as they are highly expressed in all cells. The abundance value of each protein that are pulled down by either EdU or EBndG were divided by the abundance value of beads control to attain the fold change (FC). To permit calculation of LFQ ratios for proteins with missing values in the bead controls, an arbitrary value (2222, chosen to be slightly smaller than the lowest empirical value) was used for imputation. The FC was further calculated as log base 2, and proteins that have FC log2 = 1 and higher are considered as the final number of proteins pulled down. EdU pull-down were compared with the EBndG pull-down proteins that were represented in Venn diagrams. The GO analysis of the unique EdU and EBndG-associated proteins were analyzed by ToppGene. The full ToppGene analysis were shown in Table S1.

The U2OS cells iPolKD samples in-gel digested as described above, replacing ABC buffer with Triethylammonium bicarbonate buffer (TEAB), and encoded with 10-plex Tandem Mass Tag (TMT) according to manufacturer’s protocol (Thermo Fisher). After quenching the TMT reagent with 5% hydroxylamine, TMT-labelled samples were pooled, lyophilized by vacuum centrifugation, and fractionated by C18 Reversed Phase chromatography in spin column format at pH 9. Eluates were acidified with 0.1% TFA, lyophilized, and reconstituted in 30 μl 0.1% aqueous formate. 3 μl of each fraction was analyzed in duplicate by LC-MS. Labelled peptides were resolved using an easyLC1200 nano-chromatograph (Thermo Scientific), peptides were loaded on a 25 cm x 75 μm ID column packed in house with Reprosil C18 silica particles (1.9 μm diameter, Dr. Maischt), and resolved on a 5-35% acetonitrile gradient in water (0.1% formate) at 200 nl/min. Eluting peptides were ionized by electrospray (2200 V) and transferred into a Thermo Exploris Orbitrap mass spectrometer whose ion transfer tube was maintained at 275 °C. The MS was set to collect 120,000 resolution orbitrap precursor scans (profile mode) every 3 s (m/z 380-2000 Da, AGC= 400,000 ions, maximum injection time = 50 ms, RF lens = 40%). Precursor ions were selected for data-dependent acquisition in cycle-time mode (i.e. performing as many MS2 as possible within the 3 s interval between consecutive MS1 scans), applying the following filters: dynamic exclusion 30 s with 10 ppm mass tolerance, monoisotopic ion selection based on isotopic pattern expected for peptides, charge state 2-5 (ions with undetermined charge state were excluded), and minimum ion intensity 1,000.

Precursor ions were selected using the quadrupole (isolation window = 1.2 Da), accumulated for up to 110 ms (AGC= 30,000 ions), and fragmented by HCD at stepped 26,32,38% normalized energy. Spectra were recorded in centroid mode using the orbitrap at 60,000 resolution, with first mass locked to 100 Da.

Peptide-spectral matching and reporter ion intensity extraction were performed using Proteome Discoverer v2.4, with Mascot as scoring function set as above except for MS2 mass tolerance set to 20 ppm, and TMT10-plex adduct set as fixed modification. The Abundance of each protein identified in "Pulse" and ’Wash-Chase" samples were normalized with "Beads Control". The "Pulse"/"Wash-Chase" proteins with peptide count 2 and above were used for ToppGene analysis.

### Immunofluorescent staining and click fluorophore labelling

Cells were grown overnight on glass coverslips coated with poly-D lysine. After 2 h treatment with either DMSO or variable concentration of BPDE diluted with DMSO (0.1 to 2μM), fresh media was added with either 5-EU (1 mM for 1 h) ^53^, EdU (20 μM for 10 min) or EBndG (100 μM for 30 min) and the control with DMSO for 30 mins. The cells were gently washed 2x with PBS and fixed with 4% buffered paraformaldehyde at room temperature (RT) for 20 min. Cells were washed twice with PBS containing 1% BSA or kept in 1% BSA overnight at 4°C, then permeabilized with 0.5% (v/v) Triton X-100 in PBS for 15 min. Cells were washed with PBS and click fluorophore labelling was performed by incubating with click reaction 1 mM CuSO4, 2 mM THPTA, 10 mM sodium ascorbate, 0.2 μM Alexa Fluor 594 azide (Thermofisher A10270) complemented with PBS for 1.5 h at RT in dark. The cells were washed 3x with PBS and incubated in blocking buffer containing 10% donkey serum (JacksonImmunoResearch 017-000-121) in PBS for 1 hour, washed with PBS and incubated with primary antibody solution in 1% blocking buffer overnight at 4C. Cells were then washed 3x with PBS and incubated with secondary antibody solution, consisting of 1: 2000 (v/v) donkey anti-mouse Alexa Fluor 647 (A-31571) or donkey anti-rabbit Alexa Fluor 488 (A-21206) for 2 h at room temperature. Washed 3x with PBS followed by 5 min incubation in 4′,6-diamidino-2-phenylindole DAPI (1:2000) and mounted with Vectrashield mounting medium. The following antibodies and dilutions were used: anti-POLK or anti-DinB (sc-166667, IF 1:100, WB 1:1000), anti-POLH (sc-17770, IF 1:100), anti-POLI (sc-101026, IF 1:100), REV1 (sc-393022, IF 1:100), anti-PARP1(gift from George Lucian Moldovan lab, Ab-95425 IF1:500), anti-NPM1 (HPA011384 Sigma, IF 1:250, WB 1:2500), anti-UBF1 (sc-13125, IF1:100), anti-FBL (ab137422 IF 1:250), anti-γH2AX (sc-517348, IF 1:150), anti-GAPDH (Invitrogen 39-8600 WB 1:2500), anti-vinculin (sc-73614, 1:1000).

### Isolation of Polκ synthesized DNA followed by qPCR of rDNA regions

Cells (5 × 10^6^) were treated with 0 or 0.5 μM BPDE for 2h followed by 100 μM EBndG for 30 min. Cells were pelleted and the DNA isolated by two different methods.

*Method A*. Chromatin was isolated and treated with the click reagent as described in the Isolation of protein on nascent Polκ synthesized DNA (iPolKD) section, except in the final step DNA was eluted by incubating the beads in 10 mM Tris (pH 8), 1 mM EDTA containing 1% SDS at 95 °C for 10 min. The protein was removed by two phenol/CHCl_3_/*iso*amyl alcohol (25/24/1) extractions followed by isopropanol precipitation and a 70% ethanol wash step. The DNA was redissolved in 0.5 mL 10 mM Tris-HCl (pH 8.0), 1 mM EDTA and treated with 20 μL RNase A (1mg/mL) for 30 min at 37 °C. The aqueous solution was extracted twice with 0.5 mL phenol/CHCl_3_/*iso*amyl alcohol (25/24/1) followed by isopropanol precipitation and a 70% ethanol wash.

#### Method B

5x10^6^ cells were lysed with 0.5 mL 10 mM Tris-HCl (pH 8.0), 150 mM NaCl, 1 mM EDTA, 0.5 %SDS and 20 μL proteinase K (20 mg/mL) for 3 h at 55 °C. The protein was removed by two phenol/CHCl_3_/isoamyl alcohol (25/24/1) extractions followed by isopropanol precipitation and a 70% ethanol wash step. The DNA was redissolved in 0.5 mL 10 mM Tris-HCl (pH 8.0), 1 mM EDTA and treated with 20 μL RNase A (1mg/mL) for 30 min at 37 °C followed by 5 μL proteinase K (20 mg/mL) for 30 min at 55 °C. The aqueous solution was extracted twice with 0.5 mL CHCl_3_ followed by isopropanol precipitation and a 70% ethanol wash. The DNA was quantified with a Qubit 2 Fluorometer using the Qubit dsDNA HS Assay kit. The DNA was sheared with a Covaris M220 sonicator to approximately 100 bp.

#### DNA isolation and qPCR

The DNA was conjugated to biotin with 50 μM Biotin-PEG3-azide (Vector Labs), 1 mM CuSO_4_, 2 mM THPTA, 10 mM sodium ascorbate for 2 h. The DNA was separated from the excess biotin isolated with a DNA Clean & Concentrator-5 kit (Zymo Research). The biotin-adducted DNA was isolated by incubation with 25 μL MyOne Streptavidin C1 magnetic beads in 1 M NaCl for 1 h at RT. The beads were washed twice with 100 μL 10 mM Tris-HCl (pH 8), 1 mM EDTA, 1 M NaCl and twice with 100 μL 10 mM Tris-HCl (pH 8), 1 mM EDTA. The DNA was eluted by heating at 90 °C for 10 min followed by a rapid removal of the magnetic beads.

The primers for real-time PCR are as follows: H0, 5’-GGTATATCTTTCGCTCCGAG-3’ and 5’-GACGACAGGTCGCCAGAGGA-3’; H8, 5’-AGTCGGGTTGCTTGGGAATGC-3’ and 5’-CCCTTACGGTACTTGTTGACT-3’; and H23, 5’-CCTTCCACGAGAGTGAGAAGC-3’ and 5’-TCGACCTCCCGAAATCGTACA-3’ ^54^

### rDNA FISH

The rDNA FISH experiments were based on that described by van Sluis et al ^34,55^. The FISH probe was prepared from pUC-hrDNA-12.0 plasmid (Addgene cat no. 206320). The fluorescent probe was prepared by incubating 1 μg DNA with 60 μM FAM-dUTP (Lumiprobe, cat no 2077) 10 μM dTTP:, 40 μM each of dATP, dCTP and dGTP, 0.3 U/μL DNA polymerase I (NEB) and 1 ng/mL DNase I (NEB) in 50 mM Tris-HCl, pH 7.8, 5 mM MgCl_2_, 1 mM DTT, 50 μg/mL BSA for 2h at 15°C. The reaction was quenched with EDTA and the DNA purified with a PCR purification column, with the product eluted in 10 μM Tris-HCl pH 8, 0.1 mM EDTA. The labeled probe (250 ng) was mixed with 25 μL human Cot-1 DNA (1 mg/mL) and 5 μL tRNA (10 mg/mL) then precipitated by the addition of sodium acetate and ethanol. The DNA was redissolved in 125 μL formamide.

The treated U2OS cells on slides that were conjugated with AlexaFluor594-picolyl azide, as described above, and were dehydrated by serial 5 min incubations in 70%, 80%, 90%, 100% ice-cold ethanol. After air-drying, the slides were denatured in 2X SSC containing 70% formamide at 72°C for 5 min. The slides were then dehydrated in increasing concentrations of ethanol as describe above. The slides were air-dried and then put on a 42 °C heating block for 30s to completely dry. 2× Hybrisol VII (12.5 μL, MP Biomedicals) was placed on cells on the microscope slide followed by 12.5 μL of probe (in formamide), which was denatured at 72°C for 10 min. A coverslip was added, sealed with rubber cement, and incubated overnight at 37°C. The coverslip was removed, washed twice in 2X SSC containing 70% formamide for 10 min, 2X SSC for 10 min, then in 4X SSC + 0.1% Tween for 10 minutes. The cells were examined as described above.

### RNase A, RNase H and DNase I digestion

U2OS cells grown on coverslips overnight were treated either with DMSO or 2 μM BPDE for 2 h, then incubated with fresh media with only DMSO (no nucleotide control) or 100 μM EBndG for 30 minutes. After washing with PBS, cells were incubated for 15 min either with PBS (control) or DNase I (10U) (NEB M0303) or RNase A (2mg/mL) (NEB T3018) or RNase H (NEB M0297) at 37C, fixed in 2% paraformaldehyde for 20 minutes and incubated in blocking buffer (1% BSA in PBS). Cells were permeabilized and click reaction performed for 1.5 h as described above. Cells were again incubated with either PBS (control) or DNase I or RNase A or RNase H for 15 minutes, followed by two more washes with PBS buffer. Cells were further processed for immunofluorescence as described above.

### Confocal Microscopy, Image Analysis, Quantification and Statistical analysis

Imaging was performed by super resolution confocal microscopy using Zeiss LSM910 with AiryScan2 in multiplex mode SR-2Y using a 20X (NA 0.8), 40X (NA 1.2) water and 63X oil (NA 1.4) objectives. Imaging parameters such as laser power, digital gain, digital offset, scan speed was kept constant for the same experimental batch of treated and untreated cells.

Resulting images were quantified through a custom image analysis pipeline using CellProfiler version 4.2.1 ^56^. Briefly, individual imaging channels were first split for object detection. In all images DAPI channel was used for nuclei object detection. The nuclei objects were further used as masks on EBndG, EdU, EU, PolK, UBF1 and γ-H2AX channels using the image mask function to isolate and measure the intensity within the individual nuclear object boundaries.

The fluorescence intensities of EBndG, EdU, EU, Polκ, UBF1 and γH2AX were exported as spreadsheet for statistical analysis using GraphPad Prism version 10 software. In box plot graphs, boxes represent the 25–75 percentile range with median, and whiskers represent the 5–95 percentile range. Statistical tests were applied as described in the Figure legends and were calculated with Prism 9. In all quantitative immunofluorescence studies, random cells per group were scored and mentioned in the figure legends.

### RNA transfection and Real-time PCR

Gene knockdown was performed using Lipofectamine RNAiMAX transfection reagent according to the manufacturer’s protocol. The cells were harvested after 72 h for RNA isolation using Zymo Research RNA isolation kit. cDNA prepared using SuperScriptIII Reverse

Transcriptase kit (Thermo Fisher) and OligodT primers. To check for the transcript level of pre-rRNA the cDNA was synthesized using random hexamer primers. Realtime PCR was performed using 10 ng of cDNA with SYBR Green Master mix. Samples were analyzed on a LightCycler480 and quantified with LightCycler480 quantification software. The fold change calculated by delta-delta Ct method with GAPDH as the housekeeping gene control. The siRNA used were siPOLK (pool of 3 siRNAs sc-60537), siScrambled (santa cruz), pool of 3 custom made Dicer-Substrate Short Interfering RNAs (DsiRNAs) from IDT (hs.Ri.POLI.13, hs.Ri.POLH.13, hs.Ri.REV1.13).

The primers used are: GAPDH 5’-CGACCACTTTGTCAAGCTCA-3’ and 5’-AGGGGAGATTCAGTGTGGTG-3’; 47S pre-rRNA 5’-TGTCAGGCGTTCTCGTCTC-3’ and 5-’GAGAGCACGACGTCACCAC-3’; POLK ^77^ 5’-TGAGGGACAATCCAGAATTGAAG -3’ and 5’-CTGCACGAACACCAAATCTCC-3’; POLI 5’-ACTTTCTGCGGTGACTGTGT-3’ and5’- TACATGGCTTCCCGCATCTC-3’;CDKN2A 5’-AGGAGGACGAGAGAGAGAGG-3’ and 5’- TGAGGAGGAGGAAGAGGAGG-3’ ;ACTB 5’-CACCATTGGCAATGAGCGGTTCC-3’ and 5’- GTAGTTTCGTGGATGCCACAG-3’;POLQ 5’GTCTGAGACTATTTACCAGGAACC-3’ and 5’- CCTTCCAAGGAAAAGTCTCCCTC-3’ ;BRD4 5’-GTGGGAGGAAAGAAACAGGGACA-3’ and5’-AGGAGGAGGATTCGGCTGAGG-3’; POLB 5’-CGCGTCAGTTGGGACTTAAG-3’ and 5’-ACTATCGCCAACCTCAATGC-3’; PRIMPOL 5’-TGTGGCTTTGGAGGTTACTGA-3’ and 5’- TTCTACTGAAGTGCCGATACTGT-3’; REV3L 5’-GTCTGAGACTATTTACCAGGAACC-3’ and 5’-CCTTCCAAGGAAAAGTCTCCCTC-3’; REV1 5’-GAAATCCACCGAAGAGGAGCAC-3’ and 5’-ATCAGGACTGGTCGGCAGATGT-3’; SMARCA1 5’-AACCTGGCAAGTGCTGATGTGG-3’ and 5’-CAGTGTTGTCAGTGATGAGACGG-3’; SMARCA4 5’- TACCCTCAGGACAACATGCACC-3′ and 5’-ACCGCATCCCCATTCCTTT-3’.

### Cell viability assay

Cell viability was determined using the CellTiter-Glo® 2.0 Luminescent Cell Viability Assay. Briefly, U20S cells were seeded in a 96-well plate (2,000 cells/well) and allowed to adhere overnight. The following day, cells obtained varying concentrations of EBndG or DMSO in triplicate and were incubated for 3 days. On day 3, CellTiter-Glo® Reagent was added at half the volume of cell culture media present in each well. Plates were rocked for 2 minutes and then incubated at RT for 10 min and analyzed with a luminometer (GloMax® Navigator Luminometer). Background signal from the media and CellTiter-Glo® Reagent control was subtracted from sample values. To ensure the CellTiter-Glo® Reagent was not oversaturated with ATP in the sample wells, 1 mM ATP disodium salt was used to evaluate the luminescent value near saturation.

### SDS-PAGE and western blotting

Cells were harvested and lysed in lysis buffer (50mM Tris pH8, 20mM NaCl, 10mM MgCl2, 0.1% SDS, EDTA-free Protease Inhibitor-Roche), centrifuge for 10 min and supernatant collected. Proteins were quantified by nanodrop and boiled in Laemmli buffer. Proteins were separated with 4–20% Precast SDS–PAGE gels (Biorad #4561093), transferred to PVDF membranes and were blocked with 5% milk in TBS-T buffer. Bands were detected using various antibodies at 4°C overnight. The membranes were incubated with peroxidase-conjugated secondary antibodies for 1 hour at RT before detection using ECL system. For Polκ detection supersignal west pico PLUS chemiluminescent was used (ThermoFisher).

Images captured with ChemiDoc imaging system (Biorad), and the bands quantified using ImageJ with either GAPDH or vinculin as control. All the uncropped versions of images were showed in Supplementary Figures. Secondary antibody used with anti-DinB is m-IgGκ BP-HRP (sc-516102).

## RESULTS

### TLS Polymerase κ (Polκ) is active in the nucleolus

To identify novel cellular roles of Polκ, we employed an unbiased proteomics approach to determine the proteins associated with DNA that was replicated by Polκ in cells under normal growth conditions and after exposure to BPDE. We captured the DNA replicated by Polκ by incubating cells with the novel nucleotide analog EBndG, which we previously showed to be selectively incorporated into DNA by Polκ _10,18,19_. We developed a modified aniPOND (accelerated native isolation of proteins on nascent DNA) ^57^ procedure, which we termed iPolKD (isolation of proteins on Pol kappa synthesized DNA), to capture proteins bound to the nascent DNA synthesized by Polκ. Two different human cell lines (HEK293 and GM12878) were treated with 0, 0.1, 0.5 or 2 μM BPDE and subsequently incubated with 100 μM EBndG. We noted that cell viability was not affected upon incubation with 100 μM EBndG (Figure S1A). We selected the range of BPDE concentrations and treatment time of 1.5 h based on earlier findings and observed no apparent cytotoxicity under these conditions. A high dose of 10 μM BPDE treatment for 48 h reduced cell viability to approximately 35% and induced G1 arrest in DU145 cells ^58^. A longer time exposure (24 or 48 h) of 1 μM BPDE resulted in a 20% reduction in viability, while lower concentrations (0.1 and 0.01 μM) caused minimal cytotoxicity (<10% decrease in viability) in normal human diploid lung fibroblasts ^59^. Furthermore, treatment with 2 μM BPDE for 1 h did not immediately altered NAD⁺ levels, but a significant reduction was observed 4 h post-treatment ^60^. We selected the concentrations near the IC₅₀, which were 0.7 μM for POLK⁺/⁺ U2OS human osteosarcoma cells and 0.5 μM for POLK⁻/⁻ cells after 3 days of exposure (Figure S1B).

Using iPolKD procedure, the proteins bound to the DNA containing EBndG were identified by mass spectrometry. To compare the proteins associated with EBndG with proteins present at replication forks and replication-coupled DNA repair mechanisms, we complimented our iPolKD procedure with a traditional aniPOND experiment using BPDE and 5-ethynyl-2′-deoxyuridine (EdU) (Figure 1A).

**Figure 1.**
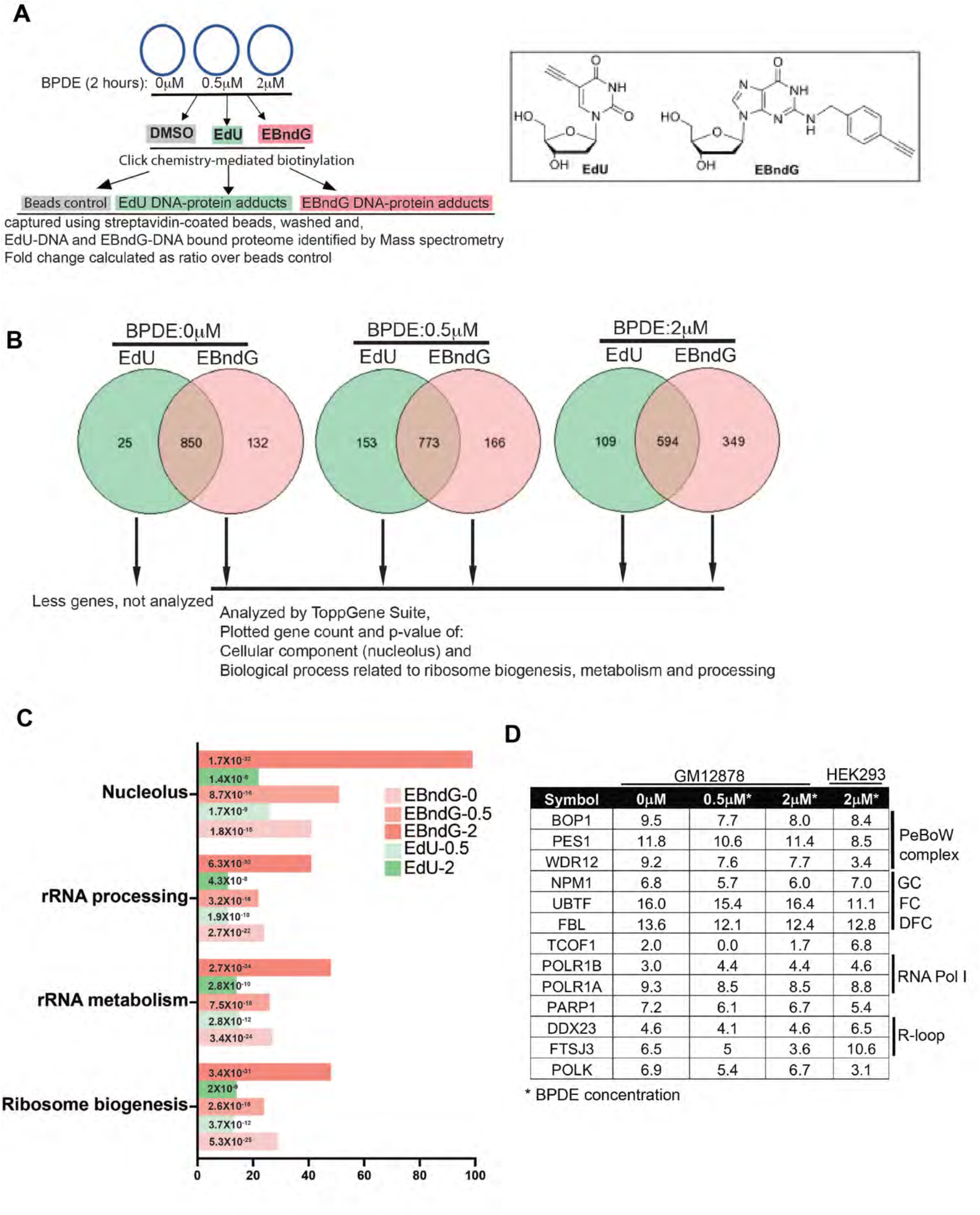
EBndG-bound proteins enriched in the nucleolus, ribosomal biogenesis and in transcriptional repression. **A.** Experimental design to identify proteins enriched at nascent DNA synthesized with Polκ captured by EBndG (iPolKD) compared to traditional aniPond. Chemical structures of EdU and EBndG shown. **B.** Venn diagrams depicting number of proteins associated with EdU (green) compared to EBndG-pull downs (red) using GM12878 cells. EdU-pull down in untreated cells identified few unique proteins (25) and were not analyzed by ToppGene Suite. **C.** GO analysis (ToppGene) of EdU and EBndG-bound unique proteins (non-overlapping protein list from venn diagrams in Fig1B analyzed). Gene counts from each pathway shown in the graph and the p-value for each pathway is indicated within each bar. EdU (green) and EBndG-pull downs (red) from cells treated with 0, 0.5 and 2 μM BPDE. See Table S1 for complete results. **D.** Representative list of proteins bound to Polκ synthesized DNA and their fold change compared to the beads control. The protein list of the EBndG-pull downs from HEK293 cells treated with 2 μM BPDE and GM12878 cells treated with (0, 0.5 and 2 μM) BPDE were analyzed with their respective beads control. Proteins grouped according to their known interacting protein complex and presence in nucleolar sub-compartments.

A Gene Ontology (GO) analysis of our data identified many DNA replication and repair proteins previously identified with EdU pull-downs ^61–63^. However, proteins associated with EBndG when compared with EdU pull-downs in both cell lines (Figure 1B, S1C) showed enrichment of nucleolar proteins required for ribosome biogenesis, processing and metabolism (Figure 1C, S1D, Table S1). Similar enrichment of nucleolar proteins was identified with Polκ synthesized DNA in U2OS cells using quantitative MS (tandem mass tag labelling) after iPolKD. Out of 511 proteins pulled-down with EBndG, 195 were nucleolar residing proteins (Figure S1E). A representation of the subset of proteins associated with nascent DNA synthesized by Polκ and their fold change compared to the beads control is shown in Figure 1D. Proteins associated with EBndG with and without BPDE damage, play roles in rDNA transcription and are present in each of the nucleolar sub-compartments. The only DNA polymerase that was enriched with the iPolKD technique was Polκ (Figure 1D). This observation supports the specificity of EBndG to Polκ and no other DNA polymerase.

### BPDE exposure induced nucleolar stress and nucleolar Polκ activity

To interrogate the novel role of Polκ in the nucleolus, super-resolution microscopy was used to locate and quantify EBndG in U2OS cells. These cells were utilized because they prominently display an average of 6 nucleoli per nucleus ^28^ and the BPDE treated S-phase arrested cells ^64^ show prominent DAPI-less nucleolar cavities. The incorporation of EBndG into DNA was visualized *in situ* by performing the copper(I)-catalyzed azide–alkyne cycloaddition (CuAAC) to covalently label the EBndG with an azide fluorophore. This approach is similar to visualizing 5-EU in nascent RNA ^53^ or EdU in nascent DNA by the click reaction. EBndG treatment resulted in low levels of incorporation throughout the nucleus with some cells showing nucleolar incorporation (0 μM BPDE panels in Figure 2A-C and S2). In cells not damaged by BPDE, nucleoli were identified by regions devoid of the DAPI signal and enriched with NPM1 (Figure 2A, C), as well as regions surrounding the FC and DFC detected by UBF1 and FBL, respectively (Figure 2D-E). With increasing BPDE exposure, nucleoli were identified by regions devoid of DAPI and NPM1 and by the formation of nucleolar caps detected by FBL and UBF1 (Figure 2D-E). Following BPDE treatment, EBndG incorporation was increased in the nucleus (Figure 2Bii, S2B-C) and became noticeably enriched in the nucleolus compared with the nucleoplasm (Figure 2C-D, S2C, S6B). This striking observation led us to explore whether BPDE treatment caused nucleolar stress and if Polκ was involved in the DNA damage stress response in the nucleolus.

**Figure 2.**
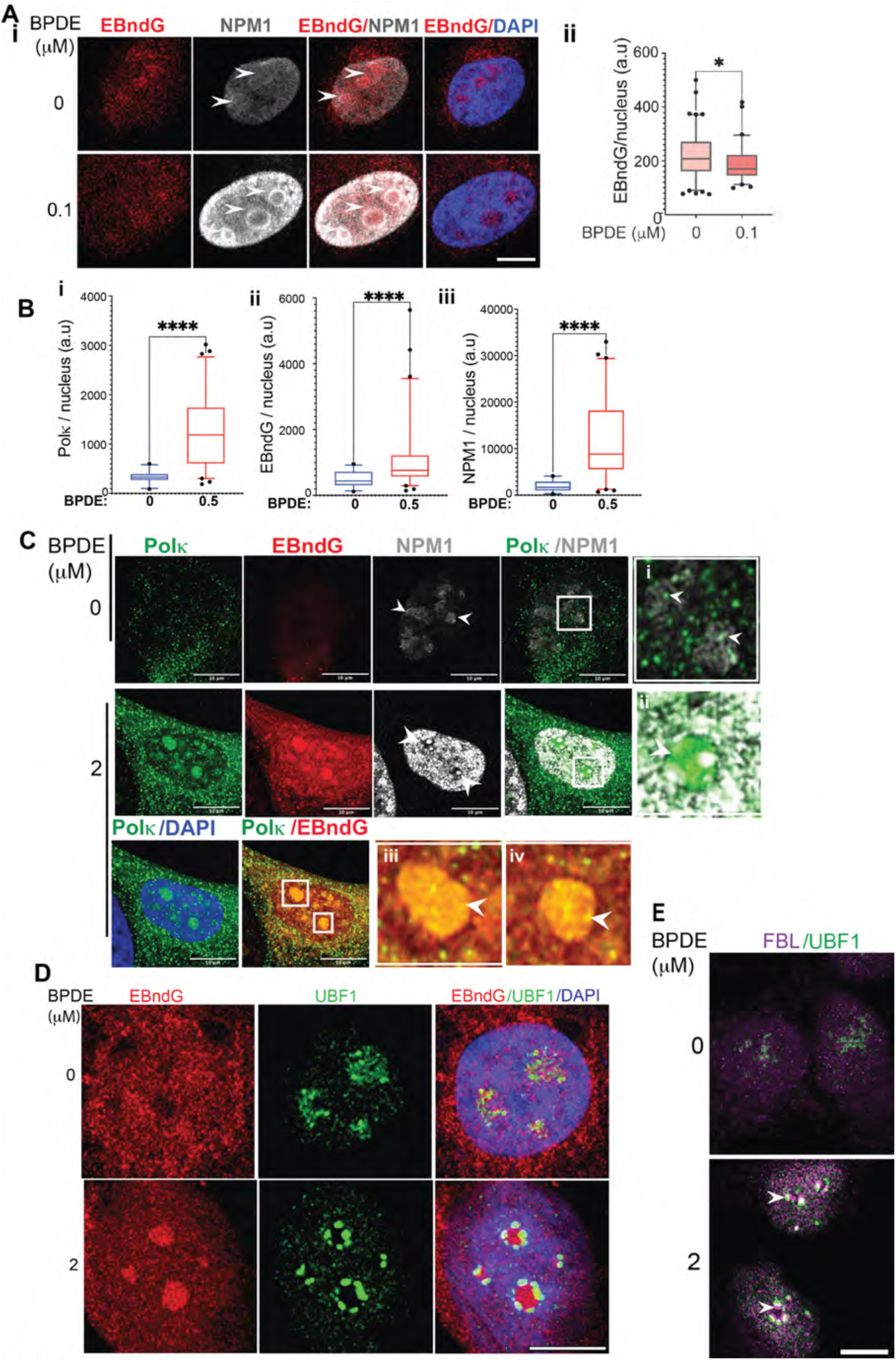
BPDE exposure induces nucleolar stress and increases the activity of Polκ in the nucleolus. **A. (i)** Representative images of U2OS cells from *N* = 2 independent experiments showing EBndG (red), NPM1 (gray) and DAPI (blue). EBndG (red) in DAPI-less regions and NPM1 in the nucleolar periphery (shown with arrows) in cells after BPDE treatment (0.1 μM). **(ii)** Quantification of nuclear EBndG intensity in untreated (n=103) and BPDE-treated (n=63) cells, represented as box plots. Boxes represent the 25–75 percentile range with median and whiskers represent the 5–95 percentile range. Data points outside of this range are shown individually. Statistical significance calculated using two-tailed Mann Whitney U test. * P = 0.0192 **B.** Quantification of total nuclear (i) Polκ, (ii) EBndG and (iii) NPM1 intensity, in 30-60 untreated and 0.5 μM BPDE-treated cells represented as box plots. Statistical analysis as Figure 2Aii. **** P <0.0001. Full images shown in Figure S2A. **C.** Representative images of untreated and BPDE-treated (2 μM) cells from *N*=3 independent experiments (quantitation in Figure S2B and full z-stack in Figure S2C). Nucleus shown by DAPI (blue), NPM1 (gray) enriched in nucleolus and nucleolar periphery in untreated cells, whereas excluded from nucleolus and present in the nucleoplasm in BPDE-treated cells. A few nucleoli are marked with arrows. Overlay panel of Polκ (green) and NPM1 shows presence of Polκ foci in the nucleoli (inset **i,ii**). DAPI and EBndG overlay with prominent EBndG enriched (red) in all DAPI-less nucleoli. Full image shown in Figure S2C. **D.** Representative images of untreated and BPDE -treated (2 μM) cells from *N* = 2 independent experiments. Untreated cell with UBF1 (green, shown with an arrow) in nucleoli and after BPDE treatment, UBF1 in the nucleolar caps (green, shown by arrows). The EBndG enriched nucleoli shown in red. Full images shown in Figure S3A. **E.** Representative cells showing FBL (magenta foci) with UBF1 (green) in nucleolar caps (shown by arrows) following BPDE treatment (2 μM). Full images shown in Figure S3B. All scale bars = 10 μm.

After treatment with various doses of BPDE followed by incubation with EBndG, we examined the localization of NPM1, as its translocation into the nucleoplasm is a typical hallmark of the nucleolar stress response ^37^. Increasing concentrations of BPDE led to an increase in NPM1 levels (Figure 2Biii, S2B, S10A), with the protein becoming concentrated on the periphery of the nucleolus (Figure 2Ai) and at higher BPDE concentrations being totally excluded from the nucleolus (Figure 2C and S2A, C). In undamaged cells, we also observed Polκ as foci throughout the cell, including in the NPM1 enriched nucleolus. However, after BPDE exposure, Polκ protein level was significantly increased (Figure 2Bi, S4D, S2B) with diffused Polκ enrichment in the nucleolus (Figure 2C and S2D-E).

We next examined the nucleolar proteins UBF1 and FBL that are present in the FC and DFC, respectively, and translocate to nucleolar caps upon nucleolar stress ^35^. A significant fraction of BPDE-treated cells contained distinct large nucleoli, with rearrangement of UBF1 and FBL proteins to nucleolar caps (Figure 2D-E, S2C). The co-visualization of EBndG and UBF1 showed EBndG incorporation into the stressed nucleoli encircled by UBF1 (Figure 2D).

Altogether, our data suggests that BPDE exposure leads to nucleolar stress.

### Nucleolar stress due to DNA damage and RNAPI inhibition induces nucleolar Polκ activity

To examine if the Polκ nucleolar activity was dependent on other DNA damaging agents, we treated U2OS cells with cisplatin and mitomycin C (MMC). Polκ has been shown to protect cells against cisplatin ^65,66^ and MMC toxicity ^67,68^, possibly in repair of interstrand crosslinks. We observed that EBndG incorporation was slightly increased in cisplatin and MMC treated cells, with enhanced accumulation in DAPI-less nucleoli (Figure 3A and S4A).

**Figure 3:**
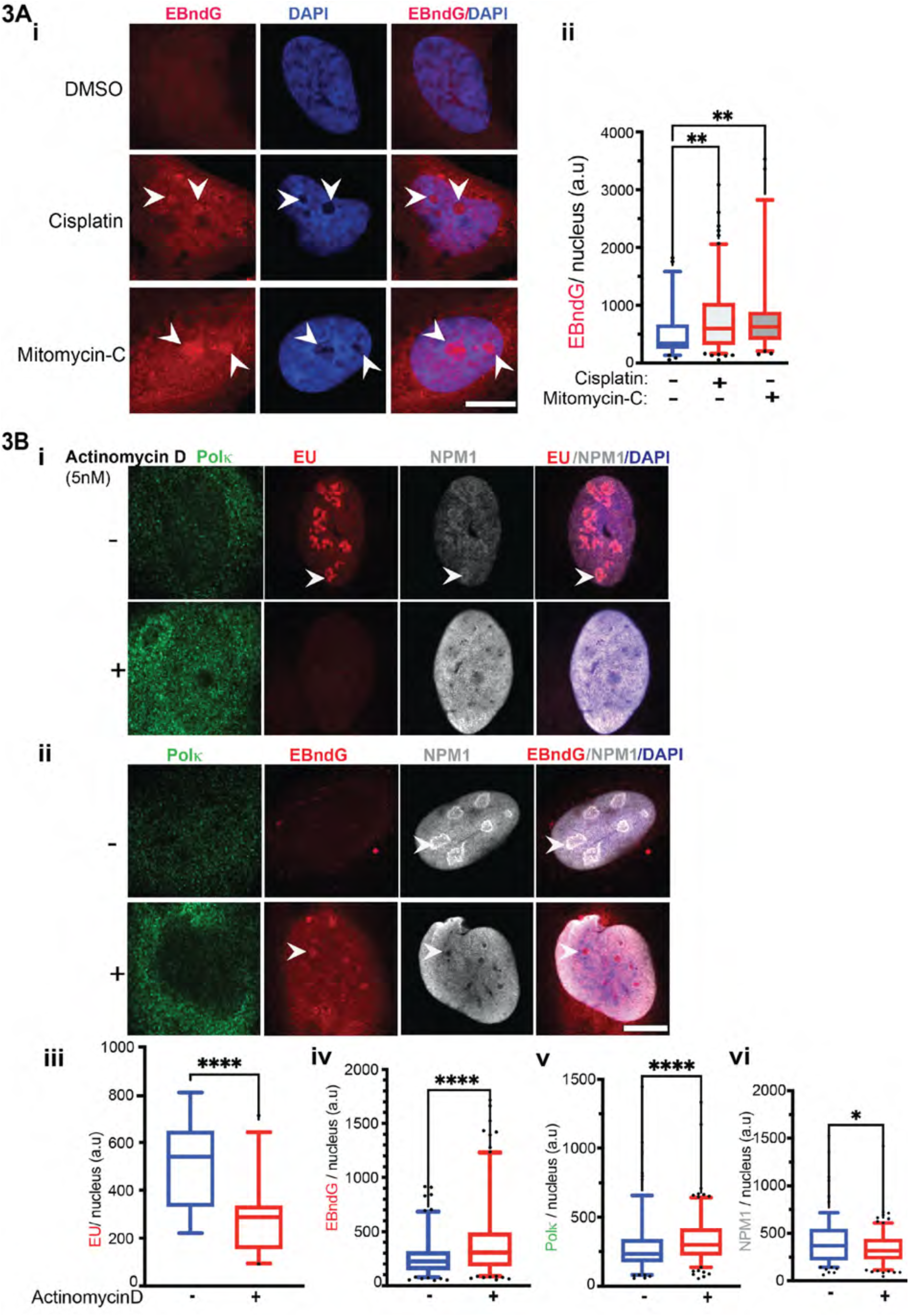
Nucleolar stress induced by cisplatin, mitomycin C and actinomycin D increases the activity of Polκ in the nucleolus. **A. (i)** Representative images of DMSO, cisplatin and mitomycin C-treated cells with EBndG (red). Overlay of DAPI (blue) and EBndG, with nucleolar enriched EBndG (red with an arrow) Full images shown in Figure S4A. **(ii)** Quantification of nuclear EBndG intensity in DMSO (n=67), cisplatin (n=125), mitomycin C (n=59) -treated cells, represented as box plots. Statistical significance calculated using Kruskal -Wallis test. *P = 0.0045. All scale bars = 10 μm. **B. (i)** Representative images of DMSO and actinomycin D-treated cells with Polκ (green), EU (red), NPM1 (gray). Overlay of DAPI (blue), NPM1 and EU shows EU in nucleolus in untreated cells (as shown by an arrow). **(ii)** Representative images of DMSO and actinomycin D-treated cells. Overlay of DAPI (blue), NPM1 and EBndG, with nucleolar enriched EBndG (red with an arrow) in actinomycin D-treated cells. Full images shown in Figure S4B. All scale bars = 10 μm. **(iii-vi)** Quantification of nuclear EU (∼25 cells), and nuclear EBndG, Polκ and NPM1 intensity in 150-160 DMSO and actinomycin D-treated cells represented as box plots. Statistical analysis as Figure 2Aii. **** P <0.0001, * P <0.0226.

Nucleolar stress is not only caused by DNA damage, but also inhibition of rRNA transcription. RNA polymerase I (RNAPI) inhibition by treatment with low-dose actinomycin D reorganizes the three nucleolar compartments, segregates the nucleolus, forms typical nucleolar caps and releases several proteins from the nucleolus ^69–73^. We questioned whether nucleolar stress caused by inhibition of RNAPI could also induce nucleolar Polκ activity. Upon actinomycin D treatment, RNAPI inhibition was demonstrated by the reduced EU incorporation in the nucleolus (Figure 3Bi, iii, S4Bi), the nucleolar stress was apparent from the NPM1 translocation from the nucleolus to the nucleoplasm and by the formation of fragmented nucleoli (Figure 3Bi, ii, S4B). Interestingly, we observed with actinomycin D treatment nuclear EBndG incorporation was significantly increased (Figure 3Bii, iv, S4B), with enrichment in DAPI-less fragmented nucleolar regions that were devoid of NPM1 signals (Figure 3Bii). This result suggests that overall nucleolar stress in cells results in enhanced Polκ activity in the nucleolus.

### Nuclear EBndG and enrichment in the nucleolus is dependent on Polκ

To confirm that the EBndG incorporation in the nucleus and the nucleolar enrichment observed after BPDE damage is dependent on Polκ, we utilized U2OS sgPolK cells that were previously shown to be deficient in Polκ _51_. We observed that U2OS sgPolK cells failed to incorporate EBndG in the nucleus or nucleolus (Figure 4Ai, Aiii, S2A). In addition, we used siRNA to knockdown Polκ in U2OS cells (siPOLK) and observed significantly reduced EBndG incorporation (Figure 4Bi, 4Biii, S4A). We observed that even after 36 h of EBndG treatment, while the siScrambled cells retained some EBndG in the nucleolus, there was not much nucleolar EBndG in the siPOLK cells (Figure S5A).

**Figure 4:**
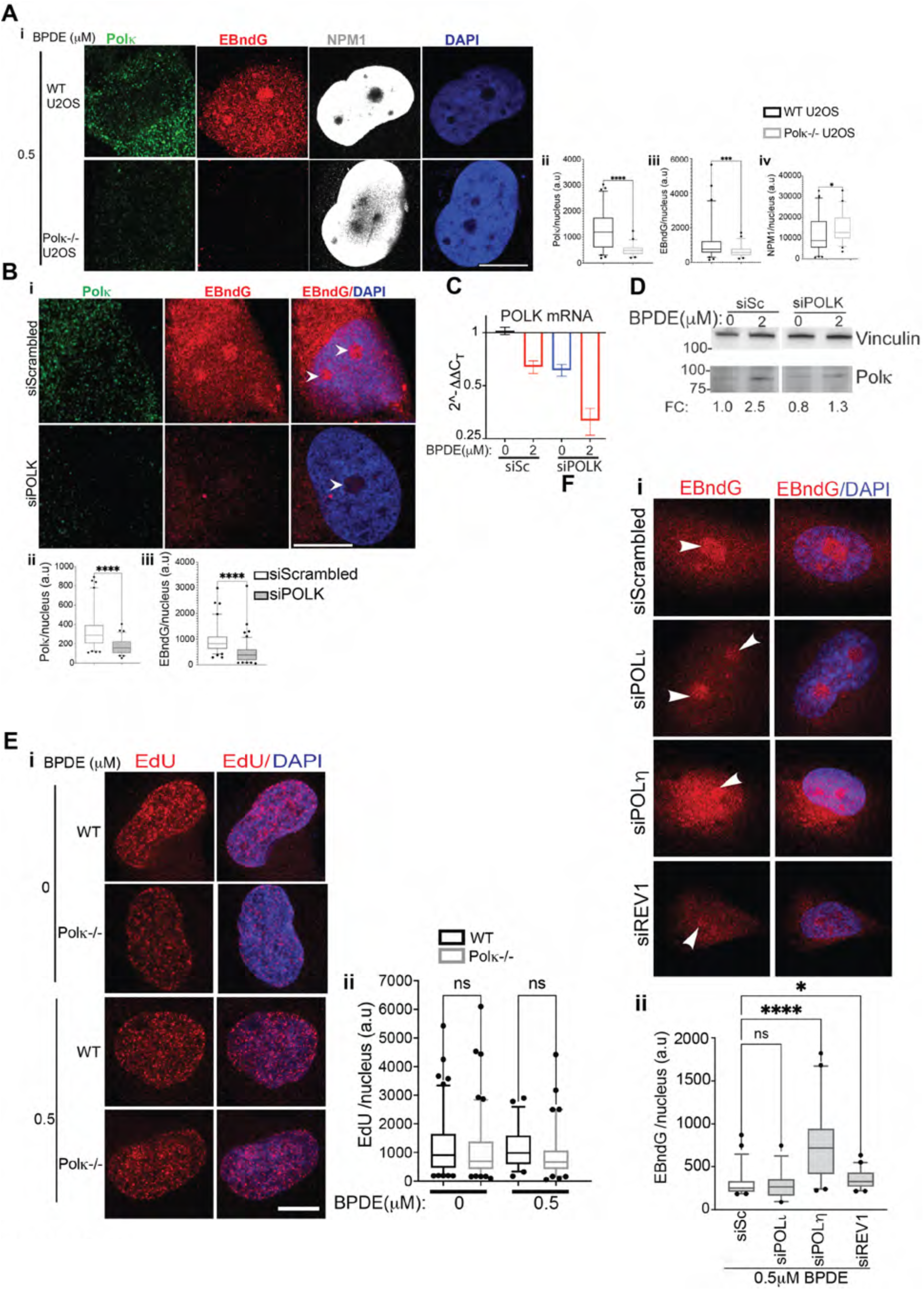
Polκ dependent incorporation of EBndG in the nucleus and nucleolus. **A. (i)** Representative images of wild type (WT) U2OS and U2OS sgPolK cells (Polκ -/-) treated with BPDE (0.5 μM). WT cells showing Polκ (green) and nucleolar enriched EBndG (red). BPDE-treated cells with NPM1 (gray) in the nucleoplasm. **(ii)** Quantification of total nuclear Polκ, NPM1and EBndG intensity in wild-type (WT) (n=63) and Polκ -/- cells (n=47), represented as box plots. Statistical analysis as 2Aii. **** P <0.0001, *** P <0.0004, * P <0.0193. **B. (i)** Representative images of cells transfected with siScrambled and siPOLK, showing EBndG (red, nucleolar EBndG shown by arrows), Polκ (green), and DAPI (blue). **(ii)** Quantification of total nuclear Polκ and EBndG intensity of siScrambled (n=80) and siPOLK (n=74) transfected cells, represented as box plots. Statistical analysis as Figure 2Aii. **** P <0.0001. Full image shown in Figure S5A. **C.** Relative expression of POLK transcript in siScrambled (siSc) and siPOLK transfected cells after treatment with 0 and 2 μM BPDE. GAPDH mRNA as internal control. Calculated with the comparative CT method (mean ± SEM). All comparisons are relative to siSc 0 μM BPDE. **D.** Western blot analysis of Polκ expression in siSc and siPOLK transfected cells, after treatment with 0 and 2 μM BPDE. Fold change (FC) of Polκ after normalization with vinculin and compared to siSc (BPDE=0). Full blots in Figure S10B. **E. (i)** EdU (red) incorporation in WT and Polκ -/- cells, after treatment with 0 and 0.5 μM BPDE, DAPI (blue). **(ii)** Quantification of nuclear EdU intensity in 0 μM BPDE-treated cells [WT (n=102) and Polκ-/- (n=103)], 0.5 μM BPDE-treated cells [WT (n=56) and Polκ -/- (n=82)]. Results represented as box plots. Statistical significance calculated using the Kruskal-Wallis ANOVA test. Full image in Figure S5C. **F. (i)** Representative images of cells and **(ii)** their nuclear EBndG intensity quantification. Cells were transfected with siScrambled (n=45), siPOLι (n=29), siPOLη (n=46), siREV1(n=49). Nucleolar EBndG (red, shown by arrows) and DAPI (gray). Statistical analysis as Figure 4Eii. **** P <0.0001, * P <0.0178. Full image in Figure S5D. All scale bars = 10 μm.

The *POLK* transcript level was reduced by ∼ 40% in siPOLK cells, and a similar decrease in *POLK* transcript level was observed upon 2 μM BPDE treatment. The combination of siPOLK and BPDE additionally decreased *POLK* transcript levels (Figure 4C).

Since BPDE damage induces transcription arrest at the sites of damage ^74^ as well as genome-wide ^75,76^, we examined the transcript levels of several genes. We found that compared to GAPDH and ActinB, transcript levels of several DNA damage response genes including *POLK* were reduced, whereas some remained unaltered or upregulated after 2 h BPDE treatment (Figure S5B). Hence, the *POLK* mRNA repression after BPDE treatment is interpreted as part of a wider transcriptional response to DNA damage ^74–77^ rather than evidence for a *POLK*-specific repression mechanism.

However, in contrast to *POLK* mRNA levels, Polκ protein levels increased ∼2-3 fold upon BPDE treatment. We observed that the steady state Polκ protein levels were inherently lower in undamaged cells, which was stabilized upon BPDE damage (Figure 2Bi, 2C, 4D, S10A).

These results suggest that BPDE treatment elevates Polκ levels by preventing its proteolytic degradation. Given that BPDE triggers the nucleoplasmic release of NPM1, which is known to stabilize Polη ^78^, it is plausible that NPM1 similarly contributes to the stabilization of Polκ under nucleolar stress conditions.

In comparison to EBndG enrichment in the nucleolus that is dependent on the presence of Polκ in the cell, EdU was incorporated as foci throughout the nucleus and was not reduced in Polκ compromised cells (Figure 4E, S5C). We also found that EBndG incorporation in the nucleolus/nucleus was not dependent on other Y-family polymerases. Cells compromised in Polη, Polι and Rev1 (transfected with respective siRNA pool) showed no reduction in EBndG incorporation (Figure 4F, S5D). In addition, we continued to observe the accumulation of EBndG in nucleolar regions. Instead, we noted an increase in nuclear EBndG incorporation in Polη compromised cells (Figure 4Fii), perhaps reflecting enhanced recruitment of Polκ to BPDE-damaged sites. This is consistent with previous studies showing that Polη’s nuclear distribution remains unchanged, however, Polκ is actively recruited to sites of DNA synthesis following BPDE treatment ^79^. Moreover, Polη-deficient XPV fibroblasts displayed a normal S-phase checkpoint response after BPDE treatment despite their inability to recover from the UV-induced S-phase checkpoint, unlike -/-MEFs ^80^, thus highlighting specifically the role of Polκ for recovery from the BPDE-induced S-phase arrest.

### Incorporation of EBndG into rDNA

Polκ was found to be highly active in regions that are GC-rich, open chromatin, and in DNA that is replicated early in the S phase by Polκ -seq ^17,18^. Since rDNA is enriched in GC-base pairs ^81^, highly repetitive, exists as open chromatin ^82–84^, replicates early in the S-phase ^85–87^, and rDNA repeats are prone to DNA damage ^24,36,36,87–89^, we queried whether EBndG was incorporated in the rDNA. We tested the presence of EBndG incorporation in rDNA using two different methods to pull-down EBndG-bound genomic DNA from either 0 or 0.5 μM BPDE treated cells (Figure 5A). In the first method, we isolated EBndG-enriched chromatin using a similar approach used in the iPolKD procedure. In the second method, we isolated genomic DNA followed by conjugation of EBndG with biotin, pulling-down biotin to release the EBndG-bound DNA (Figure S6A). The EBndG-enriched DNA was analyzed by qRT qPCR using primers that amplify the rRNA gene promoter (H0), the 28S rRNA coding region (H8), or the non-coding region (H23) ^54,90^ (Figure 5Ai). We calculated fold enrichment of EBndG pull-down rDNA loci by comparing equivalent number of cells with 100 μM EBndG versus 0 μM EBndG. We found that EBndG was incorporated in undamaged cells in all three rDNA loci. Importantly, BPDE treatment resulted in elevated EBndG insertion (Figure 5Aii, iii). In contrast, while EdU was incorporated into rDNA at higher levels than EBndG, the incorporation of EdU was decreased in BPDE-treated cells (Figure 5Aiv). The reduced incorporation of EdU into rDNA was expected as BPDE DNA damage elicits a S-phase cell-cycle checkpoint ^80^ compared to higher EdU incorporation in rDNA replication in undamaged proliferating cells. The enhanced incorporation of EBndG into rDNA in BPDE-treated cells indicates that rDNA damage induced the incorporation of EBndG into rDNA.

**Figure 5:**
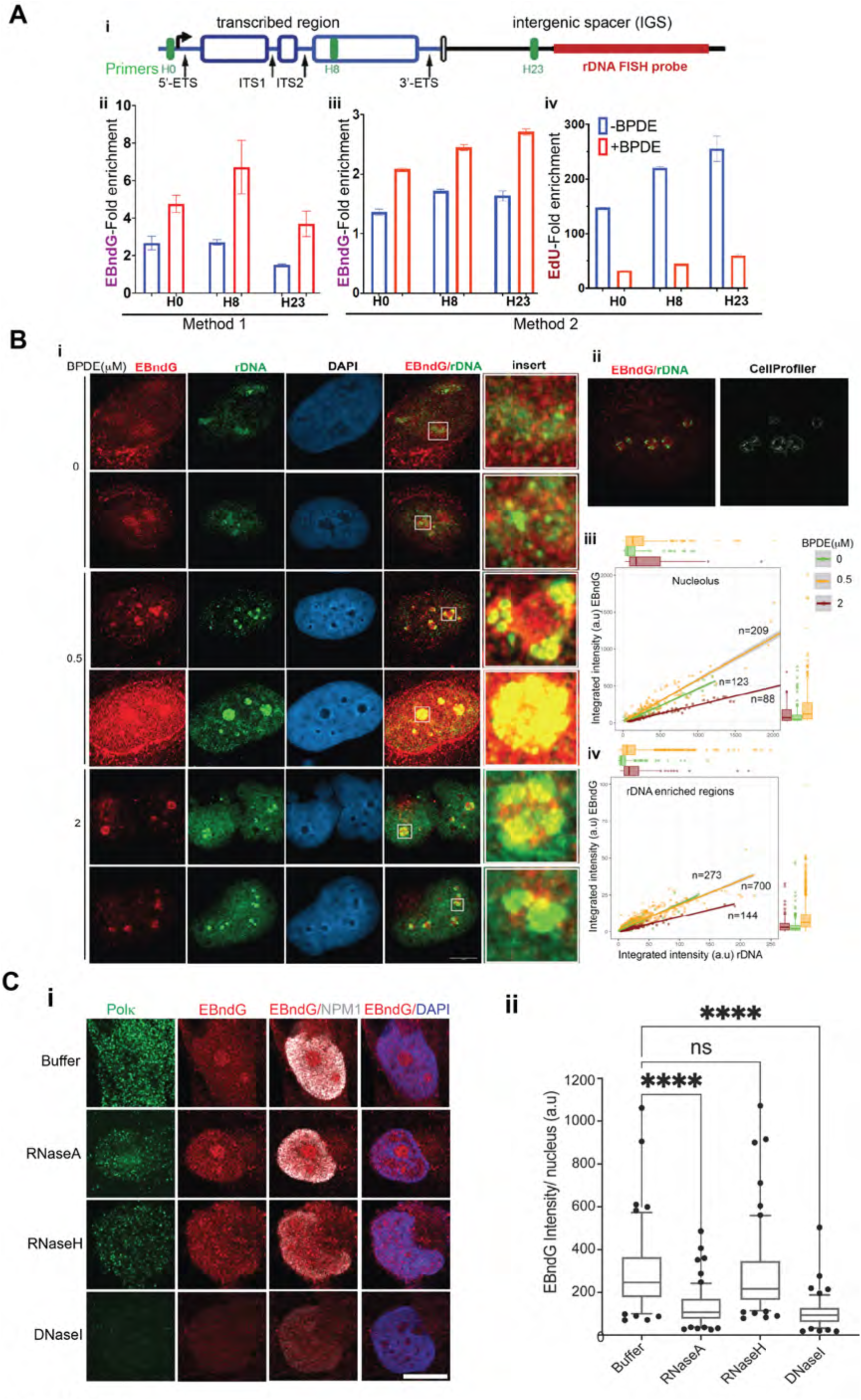
EBndG incorporation in nucleolar DNA. **A. (i)** Schematic representation of rDNA showing the transcribed region and the intergenic spacer. Also shown are the positions of the H0, H8, and H23 primer pairs (green) and the location of the FISH probe (red). Enrichment of EBndG in rDNA with 0 and 0.5 μM BPDE treatment in DNA purified by **(ii)** chromatin precipitation, and **(ii)** phenol extraction. **(iii)** Enrichment of EdU-containing DNA isolated by phenol extraction. Calculated with the comparative CT method (mean ± SEM). All comparisons are relative to beads control. **B. (i)** Representative images of cells with 0, 0.5, and 2 μM BPDE treatment. Scale bars = 10 μm. **(ii)**. Representative CellProfiler ^56^ output image in which individual nucleoli objects (in green) and rDNA intense foci (in yellow) were detected. Correlation plots of EBndG intensity versus rDNA intensity were generated for **(iii)** whole nucleoli and **(iv)** rDNA-intense foci. **C. (i)** Representative images of BPDE-treated (2 μM) cells from *N* = 2 independent experiments showing EBndG (red) enriched nucleolus, Polκ expression (green), NPM1 in the nucleoplasm (gray) and DAPI (blue) followed by either treatment with buffer, RNase A, RNase H or DNase I. **(ii)** Quantification of nuclear EBndG intensity in cells treated with buffer (n=177), RNase A (n=203), RNase H (n=236) and DNase I (n=214) represented by box plots. Statistical analysis as Figure 4Eii. **** P <0.0001. Full image in Figure S6C. Scale bar = 10 μm.

We also examined the colocalization of EBndG and rDNA using fluorescence in situ hybridization (FISH). The rDNA probe was the 12 kb fragment between 30.5 and 42.5 kb on the rDNA repeat ^55^ (Figure 5Ai). U2OS cells were treated with 0, 0.5 or 2 μM BPDE followed by incubation with EBndG. The EBndG-containing DNA was conjugated with fluorophore, and the rDNA was hybridized with the fluorescein-labelled rDNA probe. In BPDE-untreated cells, the rDNA probe identified multiple foci throughout the nucleus that would contain both active and inactive NORs (Figure 5Bi). We observed low levels of EBndG distributed throughout the nucleus, with slight enrichment in DAPI-less regions, including some colocalized with the rDNA foci (Figure 5Bi). However, following BPDE treatment, EBndG incorporation became more concentrated in the nucleolar regions as observed previously (Figure 2). Upon 0.5 μM BPDE treatment, the rDNA signal exhibited foci throughout the nucleoli in some cells, while in others it became concentrated at the nucleolar periphery, likely within nucleolar caps. At 2 μM BPDE, only large rDNA foci were observed in the nucleolus (Figure 5Bi). With BPDE treatment, we observed EBndG incorporation in nucleolar regions that both overlapped and did not overlap with the rDNA FISH signal. EBndG was observed to colocalize with rDNA foci both inside the nucleolus as well as in the nucleolar periphery in BPDE-treated cells, suggesting presence of EBndG within the nucleolar cap (Figure 5Bi). CellProfiler ^56^ was used to quantitate the spatial overlap between rDNA and EBndG signals obtained from super-resolution confocal microscopy z-stacks (63X). Individual nucleoli objects were first identified inside a nucleus and further rDNA intense foci were detected in a nested hierarchy (Figure 5Bii). Correlation plots of EBndG intensity versus rDNA intensity were generated for both whole nucleoli (Figure 5Biii) and rDNA-intense foci (Figure 5Biv). These analyses revealed a linear relationship between rDNA and EBndG signals, consistent with enhanced incorporation of EBndG within nucleoli and, more specifically, at rDNA-enriched foci. Notably, while the regression slopes for 0 and 0.5 μM BPDE-treated cells were comparable, the slope was reduced in cells treated with 2 μM BPDE. This result suggests a mechanism in which Polκ is active in rDNA regions of cells treated with 0 to 0.5 μM BPDE, but becomes less active on heavily damaged rDNA upon treatment of 2 μM BPDE.

We also compared the colocalization of rDNA with EBndG by examining the intensity of rDNA and EBndG along vectors that bisect the nucleolus (Figure S6B). In undamaged cells, we observed rDNA fluorescence in the center of the DAPI-deficient nucleolar regions. With 0.5 μM BPDE, the maximum rDNA signal is on the edge of the DAPI-less nucleoli and of the maximum of the EBndG signal. At 2 μM BPDE, the rDNA foci are at the edge of the nucleoli, but they are off the maximum EBndG signal. This analysis demonstrates that there is overlap between rDNA and EBndG incorporation, and that EBndG incorporation is enhanced in regions containing rDNA. However, EBndG incorporation is not entirely restricted to rDNA regions. In addition, there is less overlap between EBndG incorporation and more heavily damaged rDNA.

Altogether, the data suggest that although EBndG incorporation is not exclusive to damaged rDNA, overlap between EBndG and rDNA probes was consistently detected, thus demonstrating enhanced EBndG incorporation in regions containing rDNA.

Since the nucleolus is enriched with rRNA transcription and R-loops, we next tested the possibility of EBndG inserted in RNA or R-loops. We treated the cells with deoxyribonuclease I (DNase I) and observed a significant reduction in total nuclear EBndG intensity and a loss of nucleolar accumulation. Treatment with RNase H did not affect the total EBndG intensity but destroyed the nucleolar pattern that remained unaffected upon treatment with RNase A (Figure 5C, S6C). This result further suggests that EBndG is incorporated in the rDNA, possibly in regions containing R-loops.

### PCNA ubiquitination is essential for Polκ activity

To probe the mechanism by which Polκ activity is regulated in the nucleolus, we examined the role of PCNA ubiquitination that initiates the polymerase switch from a replicative to a TLS polymerase. Lysine-164 of PCNA is ubiquitinated by Rad6/Rad18 in response to replication stress induced by DNA damage or nucleotide deprivation ^91,92^. The paradigm is that ubiquitination recruits TLS polymerases to the stalled replication fork ^93–95^. PCNA ubiquitination is a critical regulator of Polκ recruitment and function for recovery from stalled forks after nucleotide deprivation ^8,51^. To examine the hypothesis that PCNA ubiquitination is important for Polκ activity in the nucleolus, we utilized 293T cells in which the endogenous PCNA gene was replaced by the K164R PCNA mutant ^50^. We observed that upon BPDE treatment, EBndG was incorporated in the 293T-WT cells and enriched in the nucleolus as expected. However, overall EBndG incorporation in the nucleus, including the nucleolus, was significantly reduced in the 293T-K164R cells (Figure 6A). This result supports the importance of PCNA ubiquitination in regulating both the nuclear and nucleolar activities of Polκ upon BPDE-induced DNA damage.

**Figure 6:**
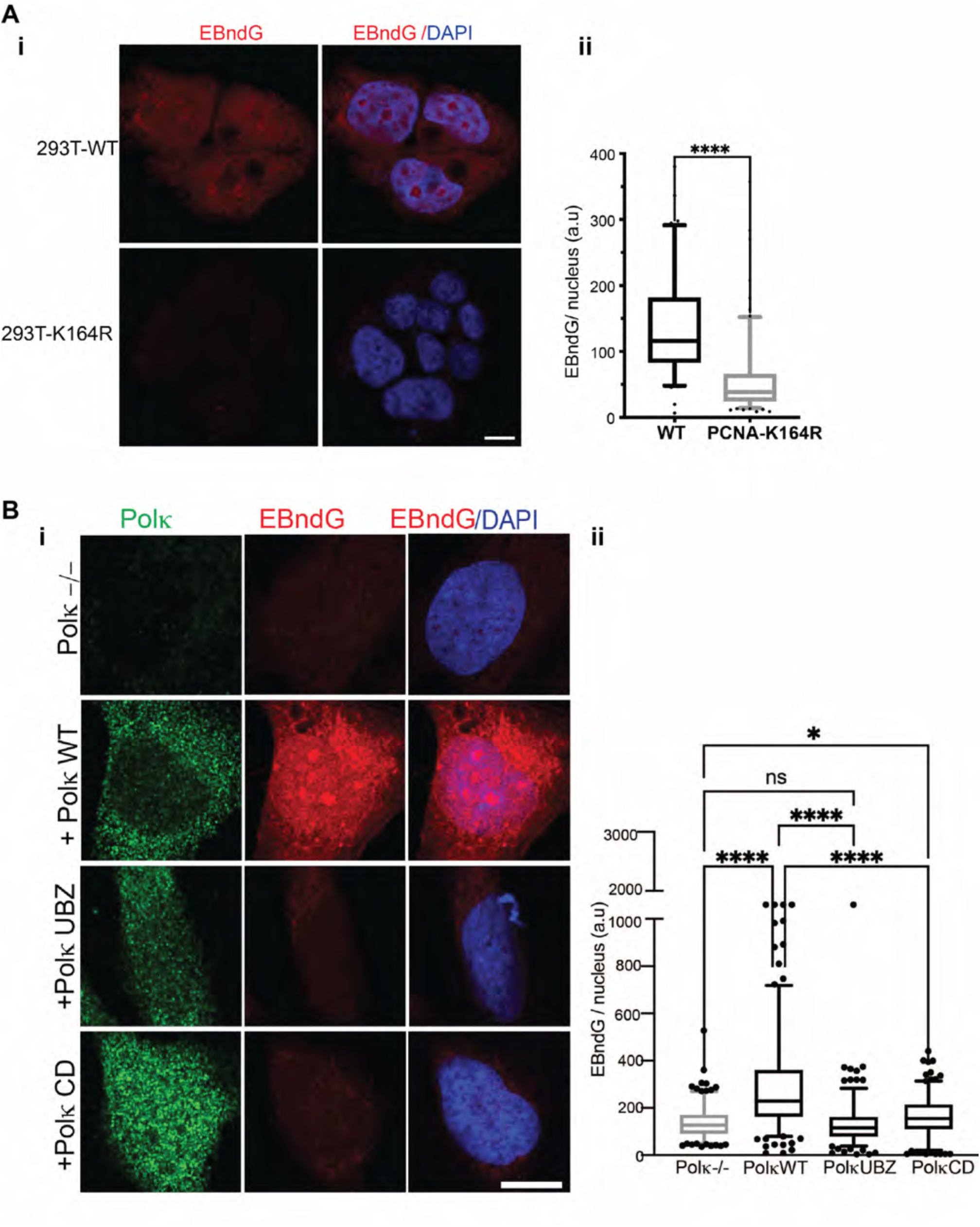
PCNA ubiquitination dependent activity of Polκ. **A. (i)** Representative images of BPDE-treated (0.5 μM) wild type (293T-WT) and PCNA K164R mutant (293T-K164R) cells. Overlay of DAPI (blue) and EBndG showing nucleolar EBndG (red) enriched nucleolus in 293-WT cells. **(ii).** Quantification of nuclear EBndG intensity of BPDE-treated 293T-WT (n=87) and 293T-K164R (n=159) cells represented by box plots. Statistical analysis as Figure 2Aii. **** P <0.0001. **B. (i)** Representative images of N=2 independent experiments. BPDE-treated (0.5 μM) U2OS sgPolK cells (Polκ -/-) and transfected with GFP-tagged PolK (+Polκ WT), GFP-tagged PolK UBZ mutant (+Polκ UBZ) or GFP-tagged PolK CD mutant (+Polκ CD) cells. Polκ expression in green, and EBndG in nucleus and nucleolar enrichment in red. Full image in Figure S7A. Scale bar = 10 μm. **(ii)** Quantification of nuclear EBndG intensity of BPDE-treated U2OS sgPolK cells (Polκ -/-, n=206) and transfected with GFP-tagged PolK (+Polκ WT, n=220), GFP-tagged PolK UBZ mutant (+Polκ UBZ, n=197or GFP-tagged PolK CD mutant (+Polκ CD, n=207) cells represented by box plots. Statistical analysis as Figure 4Eii. **** P <0.0001, * P <0.0139.

The ubiquitinated PCNA is recognized by the ubiquitin-binding zinc fingers (UBZs) present in all Y-family TLS polymerases ^96,97^. Polκ contains two UBZs ^98–100^. It was shown earlier that cells expressing GFP-tagged Polκ (PolK-WT), but not GFP-tagged Polκ with mutations in both UBZs (PolK-UBZ), interacted with ubiquitinated PCNA in cells treated with hydroxyurea ^51^. To examine whether the BPDE induced nucleolar activity of Polκ is dependent on its UBZ domain, we transiently transfected U2OS sgPolK cells with either PolK-WT or PolK-UBZ. We observed that BPDE treated U2OS sgPolK cells complemented with PolK-WT rescued EBndG incorporation with nucleolar enrichment, however, PolK-UBZ showed significantly reduced EBndG incorporation (Figure 6B, S7A). Similarly, expression of the catalytically dead mutant of PolK (PolK-CD) ^51,101^ also showed reduced EBndG incorporation (Figure 6B, S7A). We also noted that the transient expression of PolK-UBZ and PolK-CD led to fragmented nucleus lacking prominent nucleoli in ∼ 24% of cells imaged, unlike cells expressing PolK-WT that had oval nucleus with nucleolar patterns. The cells with distorted and fragmented nucleus showed overall high fluorescence without any distinguishable nucleolar pattern, hence we omitted those cells from our analysis (marked with arrow in Figure S7B). The observation of compromised activity of Polκ with either UBZ or catalytic mutations, suggested that both the DNA polymerase activity and ubiquitin-mediated interactions likely through ubiquitinated PCNA are required for BPDE-induced Polκ nucleolar activity.

### PARylation by PARP1 regulates Polκ cellular expression and activity

During DDR, one of the earliest proteins recruited is PARP1, which we found associated with EBndG on DNA newly synthesized by Polκ (Figure 1D). Approximately 40% of PARP1 molecules reside within nucleoli ^39^ and the binding of PARP1 to DNA induces its catalytic activity, synthesizing poly(ADP)-ribose (PAR) chains on itself, histone and non-histone proteins _102–104_ that recruits DDR proteins to the DNA lesions ^105^. BPDE treatment induces cellular PAR formation by PARP1 in a time and dose-dependent manner, while inhibition of PARylation by Veliparib (ABT-888) sensitizes cells to BPDE ^60^. We hypothesized that PARP1-mediated PARylation regulates the genome-wide activity of Polκ, including its function in the rDNA. We inhibited PARylation with the PARP1 inhibitor olaparib and then damaged the cells with BPDE, followed by incubation with EBndG. We found that olaparib treatment decreased EBndG incorporation in BPDE-treated cells (Figure 7A-B). We also noted that in olaparib-treated cells, Polκ protein expression was significantly reduced (Figure 7A-C). With combinative treatment of olaparib and BPDE, cells showed excessive stress, as evident by distribution of PARP1 as patches in the cytoplasm indicating that the cells were undergoing apoptosis (Figure 7A, S7).

**Figure 7:**
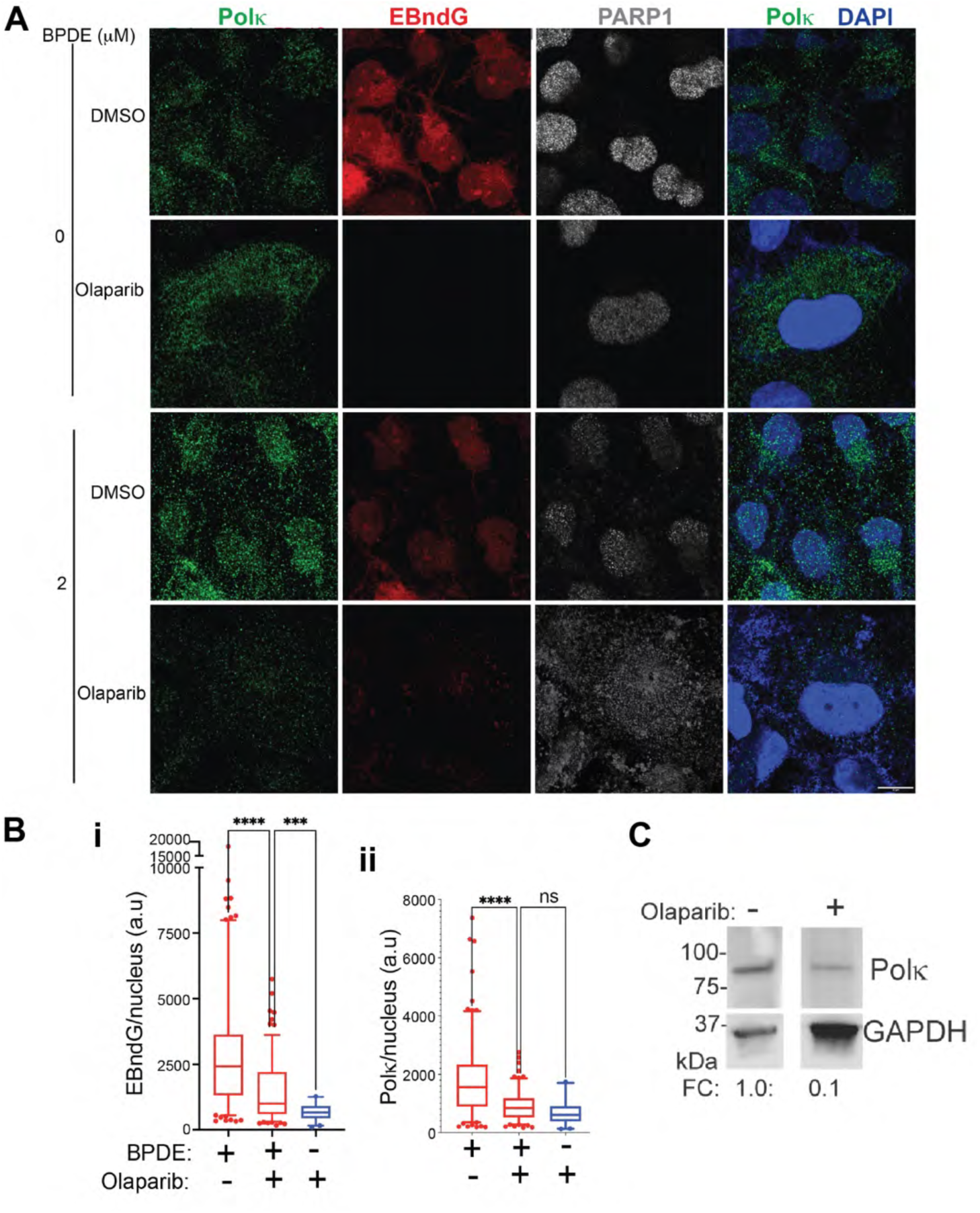
Inhibition of PARylation by PARP1 inhibitor (olaparib) decreased the stability and activity of Polκ. **A.** Representative images from *N*=2 independent experiments. Cells either treated with DMSO (control) or 10 μM Olaparib (for 3 days) (PARP1 PARylation inhibitor) were incubated with 0 or 2 μM BPDE. EBndG (red), PARP1 (gray), Polκ (green) and DAPI (blue). Full image in Figure S7A. Scale bars = 10 μm. B. Quantification of 7A, total nuclear intensity of EBndG **(i)** and Polκ **(ii)** in cells treated with olaparib and 0 μM BPDE (n=40) compared to BPDE-treated (2 μM) cells that were earlier incubated with either DMSO (n=125) or olaparib (n=153). Statistical analysis as Figure 4Eii. **** P <0.0001, *** P = 0.0002. C. Western blot analysis of Polκ expression in BPDE-treated with or without olaparib. Quantification of Polκ bands after normalization with GAPDH, FC compared to control (no olaparib treated) cells. The full blots are shown in Figure S10C.

### Polκ plays an essential role in recovering from nucleolar stress after BPDE damage

Since our experiments provide evidence that Polκ is active in the nucleolus, we next investigated whether Polκ participates in the recovery of cells from BPDE-induced rDNA damage and nucleolar stress. It was previously shown that persistent S-phase arrest in BPDE (0.1 μM) treated Polκ -/- mouse embryonic fibroblasts (MEFs) led to DSBs as detected by increased levels of γH2AX ^80^. In U2OS cells, we observed that the increase in γH2AX staining caused by BPDE-treatment was profoundly increased upon siRNA knockdown of POLK. (Figure 8A, S8A). We also noticed γH2AX foci inside and in the periphery of the nucleolus, indicating DSBs in rDNA as shown by the arrows in the +BPDE panels in Figure S8A.

**Figure 8:**
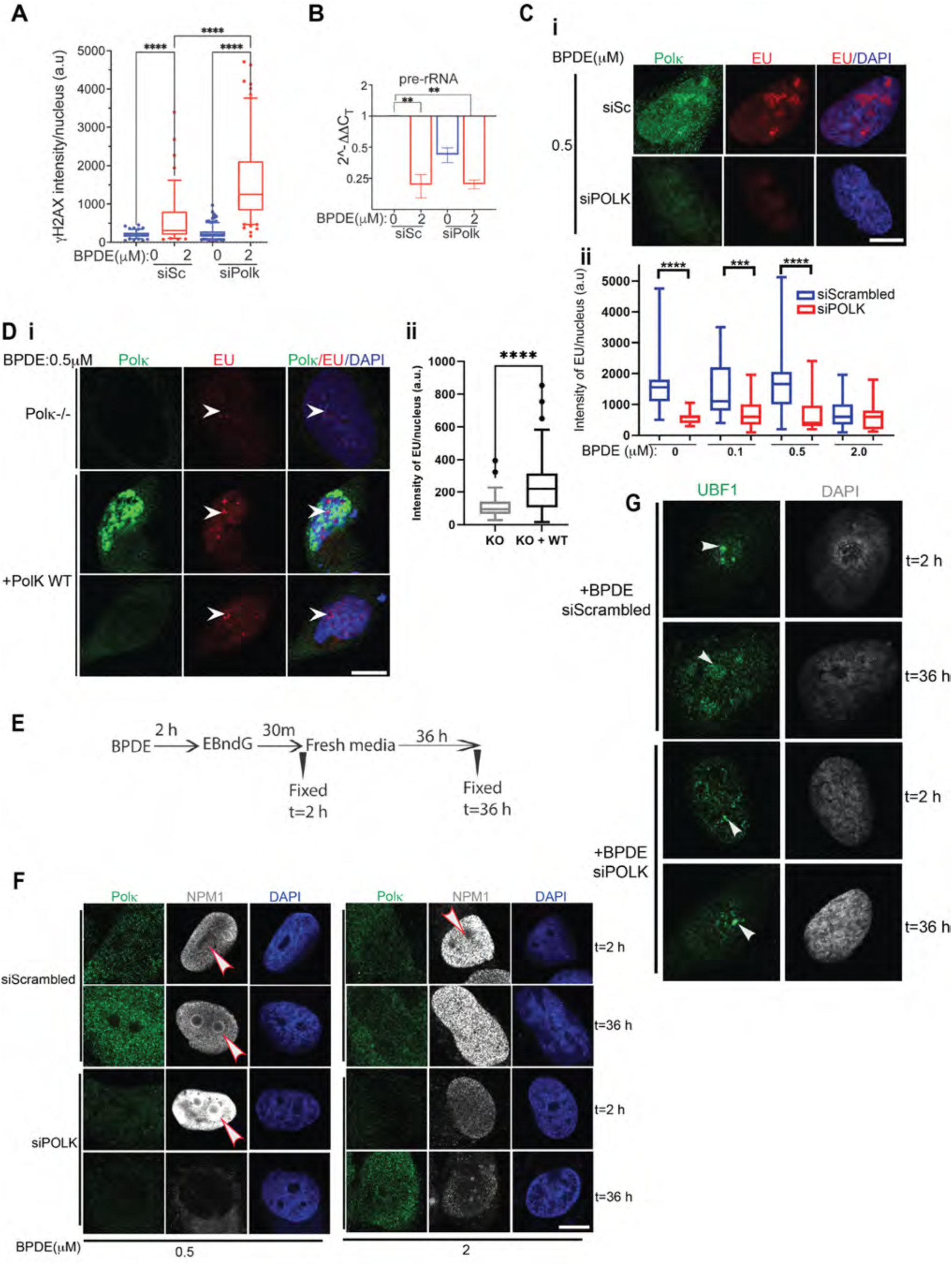
Polκ participates in recovering from nucleolar stress after BPDE damage. **A.** Quantification of nuclear γH2AX intensity in cells transfected with siSc or siPOLK after treatment with 0 and 2 μM BPDE. [siSc, BPDE 0 μM (n=136)], [siSc, BPDE 2 μM (n=96)], [siPOLK, BPDE 0 μM (n=259)], [siPOLK, BPDE 2 μM (n=141)]. Statistical analysis as described in Figure 2Aii. **** P <0.0001. Image of the cells in Figure S8A. **B.** Relative expression of 47S pre-rRNA transcript in siSc and siPOLK transfected cells, after treatment with 0 and 2 μM BPDE. GAPDH mRNA as internal control. Calculated with the comparative CT method (mean ± SEM), three technical replicates from *N*=2. All comparisons are relative to siSc 0 μM BPDE. **C. (i)** Representative images of EU incorporation (red) in nascent RNA in cells transfected with siScrambled or siPOLK. Polκ expression (green) and DAPI (blue). Full images in Figure S8B. **(ii)** Quantification of these cells (n=25-40) treated with 0, 0.1, 0.5 μM BPDE. Statistical analysis as described in Figure 2Aii. **** P <0.0001, *** P = 0.0002. **D. (i)** Representative images of BPDE-treated (0.5 μM) U2OS sgPolK cells (Polκ -/-) and transfected with GFP-tagged PolK (+Polκ WT). Polκ expression (green), nucleolar EU incorporation (red, shown by an arrow) and DAPI (blue).. **(ii)** Quantification of the nuclear EU intensity of U2OS Polκ -/- (n=35) and +Polκ WT (n=86). Statistical analysis as Figure 2Aii. **** P <0.0001. **E.** Experimental design to identify the role of Polκ in BPDE-induced nucleolar stress recovery after 36 h. **F.** Representative images of cells transfected with siScrambled or siPOLK, treated with BPDE (0.5 or 2 μM) and fixed for immunofluorescence (t=2 h) or recovered after t=36 h. Immunofluorescence showing Polκ (green), EBndG (red), NPM1 (gray) and DAPI (blue). Full images in Figure S9A. **G.** Representative images of cells transfected with siScrambled or siPOLK, treated with BPDE (0.5) and fixed for immunofluorescence (t=2 h) or recovered after t=36 h. Immunofluorescence showing UBF1 (green) and DAPI (gray). Full images in Figure S9B. All scale bars 10 μm.

Since rDNA damage will hinder rRNA transcription, we examined the effect of BPDE and siPOLK on rRNA transcription. Using RT-qPCR we detected reduced 47S primary pre-rRNA transcript in BPDE treated cells (Figure 8B). We observed that 2 μM BPDE reduced pre-rRNA transcription by factor of 4. But in addition, siRNA knockdown of Polκ reduced pre-rRNA transcription two-fold in undamaged cells (Figure 8B). We found that siPOLK did not induce a global transcriptional repression, as some genes were either unchanged or upregulated (Figure S5B). In addition, we incubated cells with 5-ethynyluridine (EU) to examine rRNA transcription in the nucleolus. As expected, we observed robust nucleolar EU incorporation in cells (Figure 8C and S8B), indicating rRNA transcription. However, EU incorporation was significantly reduced in undamaged siPOLK knockdown cells (Figure 8C and S8B). We also found that 0.1 and 0.5 μM BPDE did not eliminate EU incorporation, but that EU incorporation occurred until the cells were treated with 2 μM BPDE (Figure 8Cii). We also found that rRNA transcription was reduced in Polκ deficient U2OS sgPolK cells (Figure 8D and S8C).

Importantly, complementation of U2OS sgPolK cells with GFP-PolK WT, restored nucleolar EU incorporation, even following 0.5 μM BPDE treatment, indicating recovery of rRNA transcription (Figure 8D). These findings demonstrate that Polκ is important for repairing BPDE-induced rDNA damage and maintaining rRNA transcription

We next examined whether Polκ participates in the recovery from BPDE-induced nucleolar stress. We treated Polκ wild-type and knockdown cells with 0.5, or 2 μM BPDE for 2 h, then fixed the cells immediately (t=2 h) or allowed them to recover for 36 h in growth media (t=36 h) (schematic in Figure 8E). As observed before (Figure 2C), wild-type cells at t=2 h showed NPM1 translocated from the nucleolus to the nucleoplasm. After 36 h of recovery, NPM1 partially returned to the nucleolar periphery. However, BPDE-treated Polκ compromised cells, NPM1 redistributed throughout the cell with a majority of NPM1 in the cytoplasm (Figure 8F, S9A, S5A). Although Polκ expression in the siPOLK cells was recovering after 36 h due to the exhaustion of the siRNA, the recovery from nucleolar stress is still impaired.

To further examine the role of Polκ in recovering from nucleolar stress, we examined the formation and recovery of nucleolar caps. As illustrated in Figure 8G, treatment of 0.5 μM BPDE led to the translocation of UBF1 from the interior of the nucleoli to the periphery, an indication of nucleolar cap formation, in both siScrambled and siPOLK cells after 2 h. After 36 h, UBF1 returned to the nucleolus in siScrambled cells, while Polκ-compromised cells retained UBF1 in the nucleolar caps (Figure 8G, S9B). This result supports a role of Polκ in the recovery from BPDE-induced nucleolar stress. These recovery data further suggest that short– term (1.5 to 2 h) treatment with higher BPDE (2 μM) causes DNA damage and transient nucleolar stress but does not kill the cells. However, we observed that after a prolonged exposure of 11 days, even lower BPDE (0.5 μM) treatment significantly reduced the number of colonies. Moreover, POLK-/- cells are affected more, as observed by a reduction in the colony size and number (Figure S9C).

## DISCUSSION

We demonstrated that the Y-family TLS polymerase Polκ is active in the nucleolus. The nucleolar activity of Polκ is profoundly increased upon conditions of nucleolar stress. Both DNA damage–inducing agents (BPDE, cisplatin, and mitomycin C) and inhibition of rRNA synthesis (actinomycin D) significantly increased nucleolar Polκ activity. We also found that Polκ protected the cells from nucleolar stress, enabling maintenance of rDNA transcription low levels of BPDE exposure (Figure 8), thereby identifying a previously unrecognized role for Polκ in nucleolar homeostasis.

Leveraging the unique ability of Polκ to insert EBndG into DNA, we established a novel tool, iPolKD-MS. We found that nucleolar proteins were highly enriched on DNA that had been synthesized by Polκ (Figure 1). Using super-resolution confocal microscopy, the presence and activity of Polκ in the nucleolus was found to be increased after exposure to BPDE, cisplatin, MMC and actinomycin D (Figure 2-4). Although these agents differ mechanistically—BPDE forming bulky *N*^2^-BP-dG adducts that can lead to double-strand breaks^80,106^ (Figure 8), cisplatin and mitomycin C generating mono-adducts and interstrand crosslinks^107,108^, and actinomycin D intercalating into GC-rich DNA to inhibit RNAPI transcription^109^—all converge on inhibition of rDNA transcription and induction of nucleolar stress. ^110–112^

We showed that BPDE treatment induced the nucleolar stress response by the formation of nucleolar caps containing UBF1 and FBL (Figure 2) the release of NPM1 from the nucleolus into the nucleoplasm (Figure 2) and inhibition of RNAPI transcription of pre 47S rRNA and EU incorporation (Figure 8). Interestingly, we observed BPDE-dose dependent redistribution of damaged rDNA. At a lower dose of 0.5 μM BPDE, we observed partial intranucleolar retention and less peripheral/cap-associated localization. At a higher dose of 2 μM BPDE, we observed larger puncta that were formed either inside or at the peripheral/cap of the nucleolus (Figure 5).

Our data indicate that Polκ activity is not restricted to S phase and is instead largely cell cycle– independent, suggesting a primary role in DNA repair rather than replication. Mechanistically, Polκ activity required PARP1 activation, PCNA ubiquitination, and an intact UBZ domain, consistent with canonical TLS pathway engagement (Figures 6 and 7). While Polκ has been implicated in nucleotide excision repair (NER) ^9,10^ and interstrand crosslink repair^11,12^, and we previously estimated that ∼40% of Polκ activity reflects global genome NER ^10^, the extent to which NER contributes specifically to nucleolar Polκ activity remains unclear.

We identified Polκ as a key factor in maintaining rDNA transcription. Depletion of Polκ resulted in reduced rDNA transcription in undamaged cells, indicating a basal requirement for Polκ in nucleolar transcriptional activity. Upon exposure to BPDE, rDNA transcription was diminished in Polκ-depleted cells at 0.1 and 0.5 μM, whereas transcriptional levels were largely preserved in control cells (Figure 8). These findings suggest that Polκ contributes to sustaining rDNA transcription under conditions of DNA damage. However, at higher damage levels (2 μM BPDE), this protective effect was lost, indicating a threshold beyond which Polκ activity is insufficient (Figure 8). Consistent with these functional data, rDNA FISH analysis revealed heterogeneous nucleolar organization following 0.5 μM BPDE treatment. Specifically, some nucleoli displayed rDNA localized at the periphery, resembling nucleolar caps associated with transcriptional repression, while others retained centrally localized rDNA indicative of active transcription (Figure 5). This coexistence of transcriptionally active and inactive nucleoli is consistent with established nucleolar heterogeneity ^113–115^. Importantly, the persistence of transcriptionally active nucleoli at 0.5 μM BPDE in wild-type and siScrambled cells, as observed by both rDNA FISH and EU incorporation, supports the conclusion that Polκ is required to maintain rDNA transcription in the presence of DNA damage.

Incubation of EBndG-treated cells with DNase I and RNase A indicated that the site of EBndG incorporation is DNA. The loss of the nucleolar signal with RNase H treatment suggests an association with R-loops in which hydrolysis of the RNA allowed the DNA-bound EBndG to diffuse throughout the nucleus (Figure 5). R-loops are transient, reversible structures that form in many regions of the genome. Predictors of R-loop presence include high transcriptional activity, open chromatin and high GC content, all present in rDNA ^116,117^. R-loops participate in the regulation of transcription, but they also represent a potential source of damage, and their presence must be carefully regulated ^118^. The regulation of R-loops depends on RNA-methylation, including METTL3/14 dependent N^6^-methyladenosine (m^6^A) and METTL8 dependent 3-methylcytosine formation ^54^. m^6^A RNA was shown to selectively recruit Polκ to UV-damage sites to facilitate repair ^119^ . Strikingly, independent support for a functional connection between Polκ and R-loops comes from a recent study demonstrating that Polκ physically interacts with the RNA helicase DDX23 and is enriched at R-loop loci in chromatin genome-wide; loss of either Polκ or DDX23 alone led to augmented R-loop accumulation, establishing that Polκ actively recruits DDX23 to promote R-loop resolution ^120^.

Our iPolKD-MS analysis identified DDX23 as well as other R-Loops associated proteins like pre-rRNA 2’-O-ribose RNA methyltransferase (FTSJ3) enriched at Polκ active sites (Figure 1, Table S1). FTSJ3 participates in the processing of the 34S pre-rRNA to 18S rRNA and in 40S ribosomal subunit formation ^121^, as well as a critical regulator of R-loop homeostasis ^122^.

Together, these findings converge on the conclusion that Polκ functions at or near R-loops within the nucleolus and raise the possibility that its nucleolar activity involves R-loop resolution at rDNA, a locus with exceptionally high R-loop propensity.

We also observed a disconnect between *POLK* mRNA and protein levels following BPDE exposure. BPDE damage induces transcription arrest at the sites of damage ^74^ as well as genome-wide ^75,76^; we found that the level of *POLK* mRNA was reduced after BPDE treatment (Figure 4C). However, we observed that the Polκ protein level was increased (Figure 4D, 8, 5). Given the short half-life of Polκ (∼5 h) ^123^, this suggests post-translational stabilization. NPM1 has been shown to regulate TLS by preventing proteasomal degradation of DNA Polη ^78^. It is possible that NPM1’s upregulation and translocation into the nucleoplasm in BPDE-exposed cells helped in maintaining Polκ protein stability, increasing its activity in maintaining genomic stability and thus promoting cell survival. Although Polκ was not detected in the nucleolar proteome ^43,124^, we observed increased Polκ levels in the nucleolus after BPDE damage and in the rDNA (Figure 2C and 5). Thus, we postulate that carcinogens causing nucleolar stress stabilizes Polκ protein in the cell with an increased localization in the nucleolus leading to higher nucleolar activity, specifically in the rDNA.

In summary, BPDE-induced DNA damage triggers Polκ activity that is dependent on PARP1, PCNA ubiquitination consistent with the DNA damage response (Figure 6 and 7). Sufficient rDNA damage leads to the nucleolar stress response characterized by rDNA transcriptional inhibition, nucleolar cap formation, and NPM1 relocalization (Figure 2, 8). Under these conditions, Polκ is stabilized and recruited to the nucleolus, where its activity is markedly increased. Polκ-dependent DNA synthesis supports repair of damaged rDNA and promotes recovery of rRNA transcription. The activity of Polκ was critical in maintaining rDNA transcription during low levels of DNA damage and in recovering cells from BPDE-induced nucleolar stress. Thus, in our study, we identified the novel role of Polκ in repairing damaged rDNA, resuming rRNA synthesis and recovering from nucleolar stress caused by BPDE (Figure 9). A limitation of the present study is that, although our findings support a role for Polκ in the repair of damaged rDNA and maintenance of nucleolar transcription, they do not define the relative contribution of nucleolar versus genome-wide Polκ functions to overall BPDE tolerance. In addition, while nucleolar Polκ activity is dependent on the TLS activation pathway (Figure 6), the exact mechanism of Polκ in this process will require future mechanistic studies. However, this study opens a new direction raising several questions: whether NPM1 directly regulates Polκ’s stability, and how Polκ regulates rRNA transcription.

**Figure 9:**
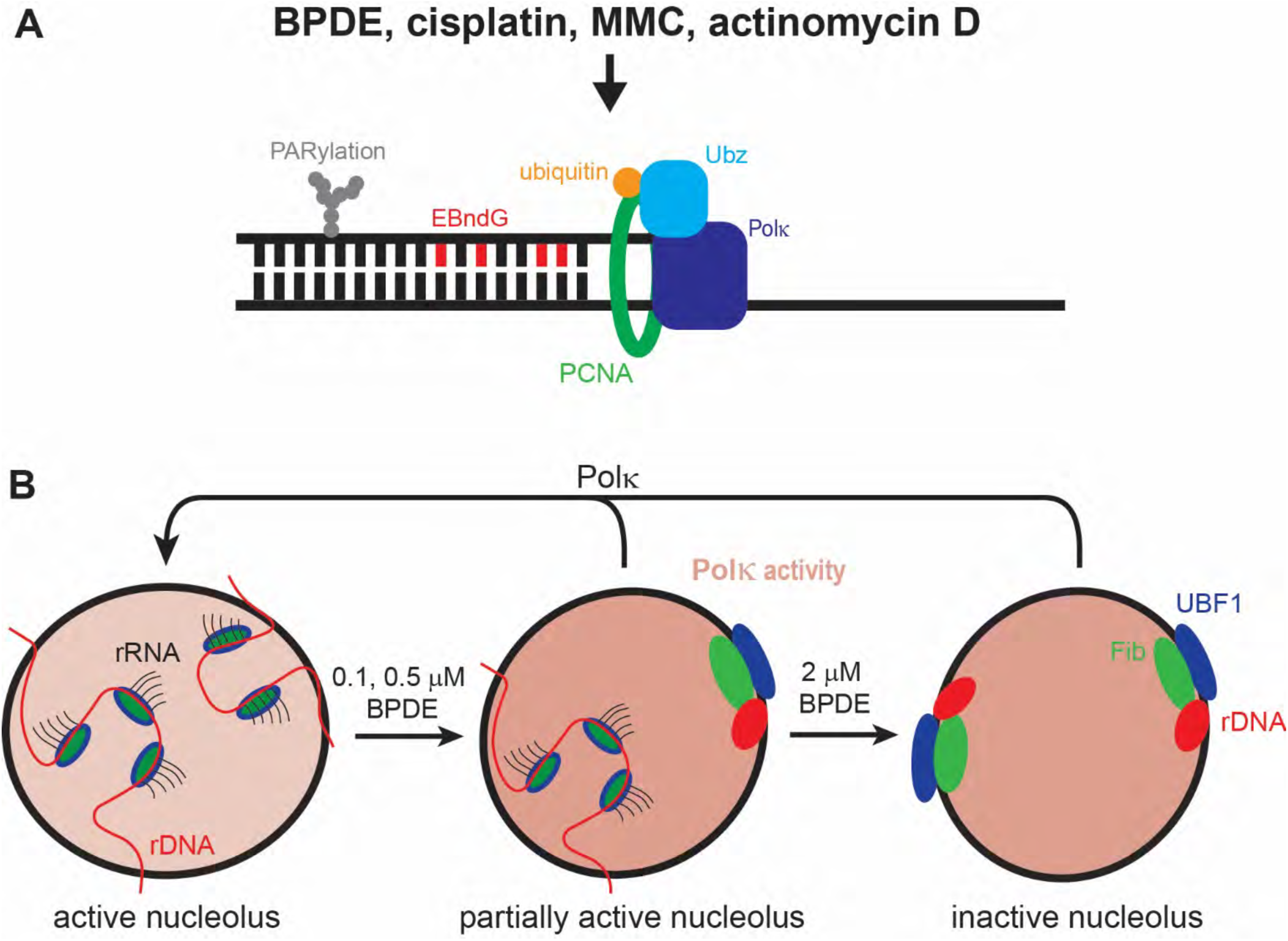
Working model on the role of Polκ in recovering from nucleolar stress after BPDE damage. A. Upon exposure to DNA damaging agents (BPDE, cisplatin, MMC) or RNAPI inhibitor (actinomycin D), DNA damages are formed on genomic DNA including rDNA. Polκ is activated by PARP1-catalyzed PARylation on chromatin, and PCNA ubiquitination that results in the incorporation of EBndG into DNA via a DNA-repair process. B. Polκ is recruited to the nucleolus and its activity in the nucleolus is evident by enrichment of EBndG in the nucleolus and rDNA. Polκ activity helps to maintain rRNA transcription in the presences of low levels of DNA damage. Upon increasing damage, nucleolar caps are formed to allow repair of DSBs. Polk facilitates cells to recover of rDNA damaged induced nucleolar stress. C. Upon exposure to DNA damaging agents (BPDE, cisplatin, MMC) or RNAPI inhibitor (actinomycin D), DNA damages are formed on genomic DNA including rDNA. Polκ is activated by PARP1-catalyzed PARylation on chromatin, and PCNA ubiquitination that results in the incorporation of EBndG into DNA via a DNA-repair process. Polκ is recruited to the nucleolus and its activity in the nucleolus is evident by enrichment of EBndG in the nucleolus and rDNA. Polκ activity helps to maintain rRNA transcription in the presences of low levels of DNA damage. Upon increasing damage, nucleolar caps are formed to allow repair of DSBs. Polk facilitates cells to recover of rDNA damaged induced nucleolar stress.

Since recent evidence connect nucleolar stress as a hallmark in a variety of cancers, neurological and aging-associated diseases, our report demonstrating the critical role of Polκ in nucleolar stress recovery and rDNA maintenance reveal new therapeutic strategies for multiple disease conditions.

## ACKNOWLEDGEMENTS

We would like to thank G. Lucian Moldovan for critically reading the manuscript, providing olaparib, the PARP1 and vinculin antibody, and the 293T-K164R cells. Benzo[*a*]pyrene diol epoxide was provided by the Organic Synthesis Core of the Penn State Cancer Institute. We would like to thank Peter Tonzi for technical assistance with sgU2OS sgPolK cells.

## AUTHOR CONTRIBUTIONS

Shilpi Paul: Conceptualization, Investigation, Data Curation, Supervision, Writing – original draft, Writing – review & editing; Abbey Rebok: Investigation; Paolo Cifani: Formal Analysis, Investigation; Writing – review and editing; Anirban Paul, Software, Formal Analysis, Methodology, Writing – review and editing; Darryl Pappin, Methodology; Tony T. Huang, Resources, Writing – review and editing; Thomas E. Spratt: Conceptualization, Resources, Supervision, Writing – original draft, Writing – review & editing.

## SUPPLEMENTARY DATA

Supplementary Data are available at NAR online.

---

**Figure S1.**
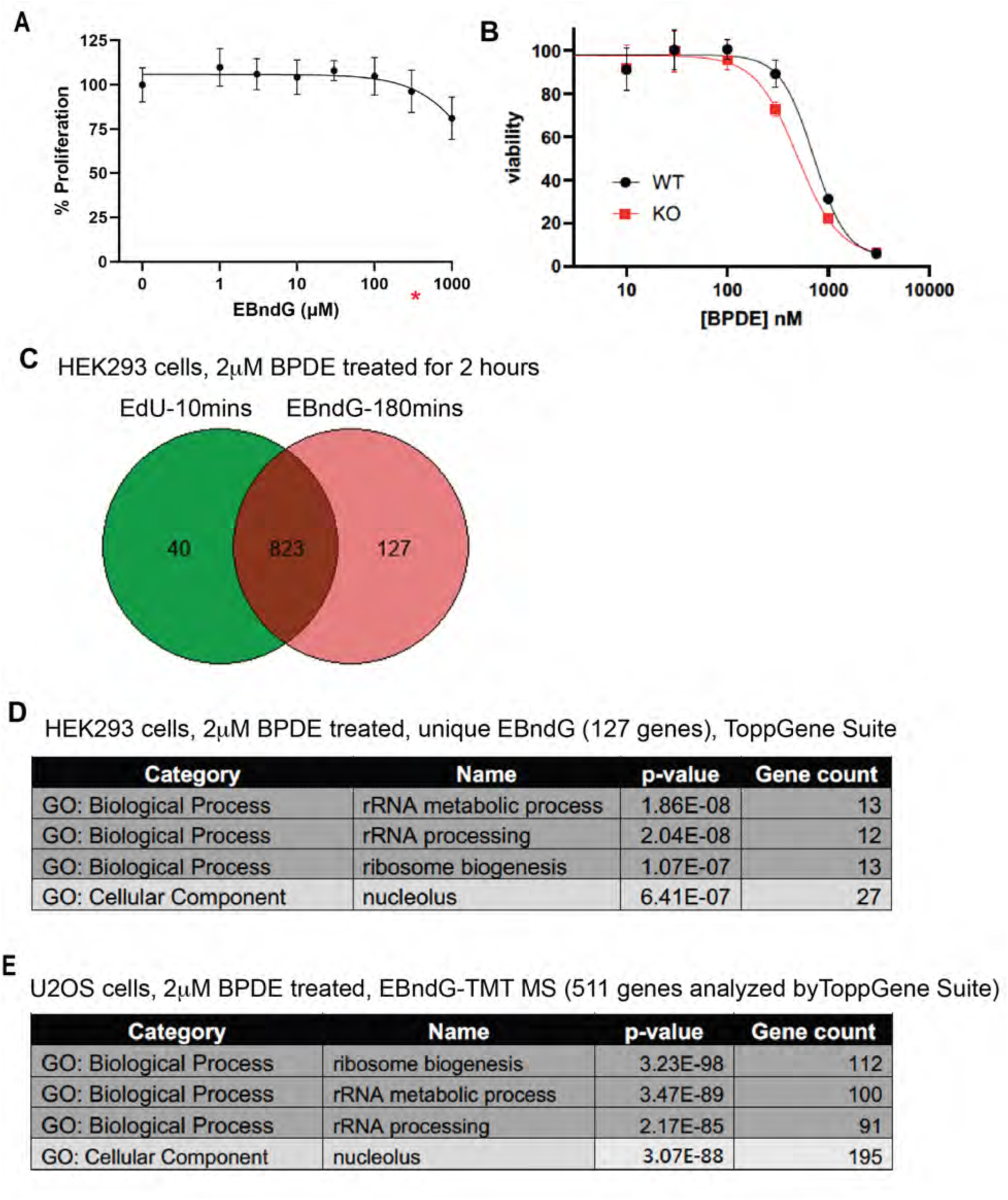
(Supplement related to Figure 1) EBndG-bound proteins enriched in the nucleolus, ribosomal biogenesis and in transcriptional repression. A. Cell viability assay of U2OS cells with increasing doses of EBndG. The working concentration of EBndG (100 μM) used in this study shown on the graph by red asterisk. B. Cellular toxicity of BPDE in U2OS cells. Cells (WT and POLK(-/-)) were treated with appropriate concentrations of BPDE in DMSO for 72 h. The viability was measured with the CellTiter-Glo Assay with 3 technical replicates. The percent viability was normalized to 100 with 0 μM BPDE. The data points are the mean ± standard deviation of 3 biological replicates. The line is the best fit to 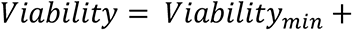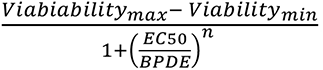, where n is the Hill slope. The fitted parameters with standard errors are: EC50 = 714 ± 68 nM and n =-2.6 ± 0.5 for wild-type cells and EC50 = 492 ± 55 nM and n =-2.0 ± 0.3 for POLK(-/-) cells. C. Venn diagram depicting number of proteins associated with EdU (in green) compared to EBndG-pull down (in red) using HEK293 cells. D. GO analysis (ToppGene) of EBndG-bound unique proteins (127). See TableS1 for complete results. E. GO analysis (ToppGene) of EBndG-bound total protein list (511) identified by iPoKD followed by TMT MS using U2OS cells treated with 2 μM BPDE. See TableS1 for complete results.

**Figure S2.**
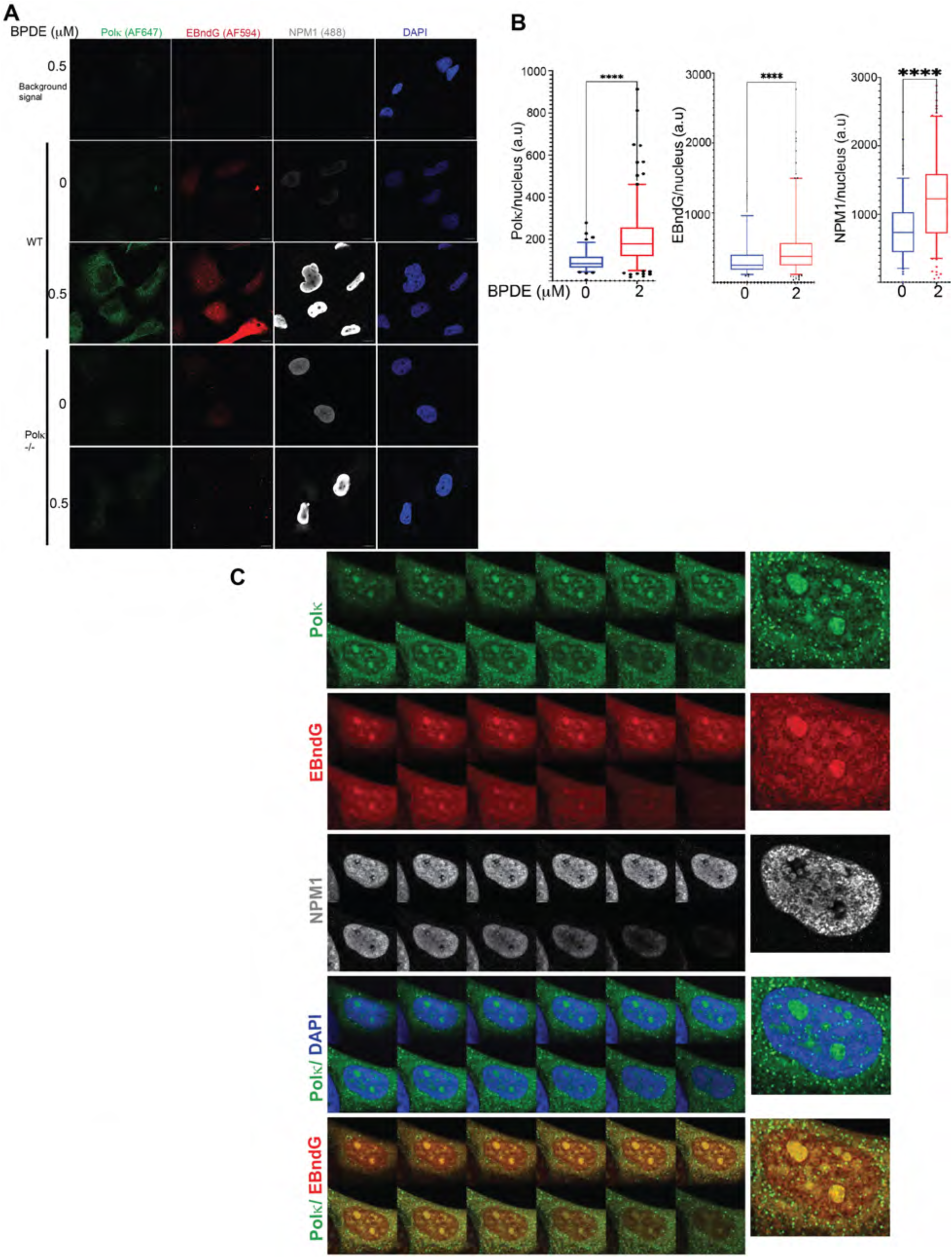
(Supplement related to Figure 2 and 4B) Nucleolar stress induced by BPDE increases the activity of Polκ in the nucleolus. **A.** Representative image panels of Fig 2B and 4B. Wild type (WT) U2OS and U2OS sgPolK cells (Polκ -/-) BPDE-treated (0 or 0.5 μM). Experiments performed for both cell lines under similar conditions. Background signal panel were cells treated under similar conditions with BPDE, without any incubation with EBndG and primary antibodies. Click reaction, immunofluorescence and imaging were performed maintaining similar conditions as experimental coverslips. AF594 panel showed a diffused red signal throughout the cells, no prominent nucleolar signal in red. Polκ shown in green, EBndG in red, NPM1 in gray and DAPI in blue. **B.** Quantification of total nuclear intensity of Polκ, EBndG and NPM1 in untreated (n=94) and 2 μM BPDE-treated (n=194) cells represented as box plots. Statistical significance calculated using Mann Whitney U test. ****P <0.0001. **C.** Full z-stack image panels of Fig 2C.

**Figure S3.**
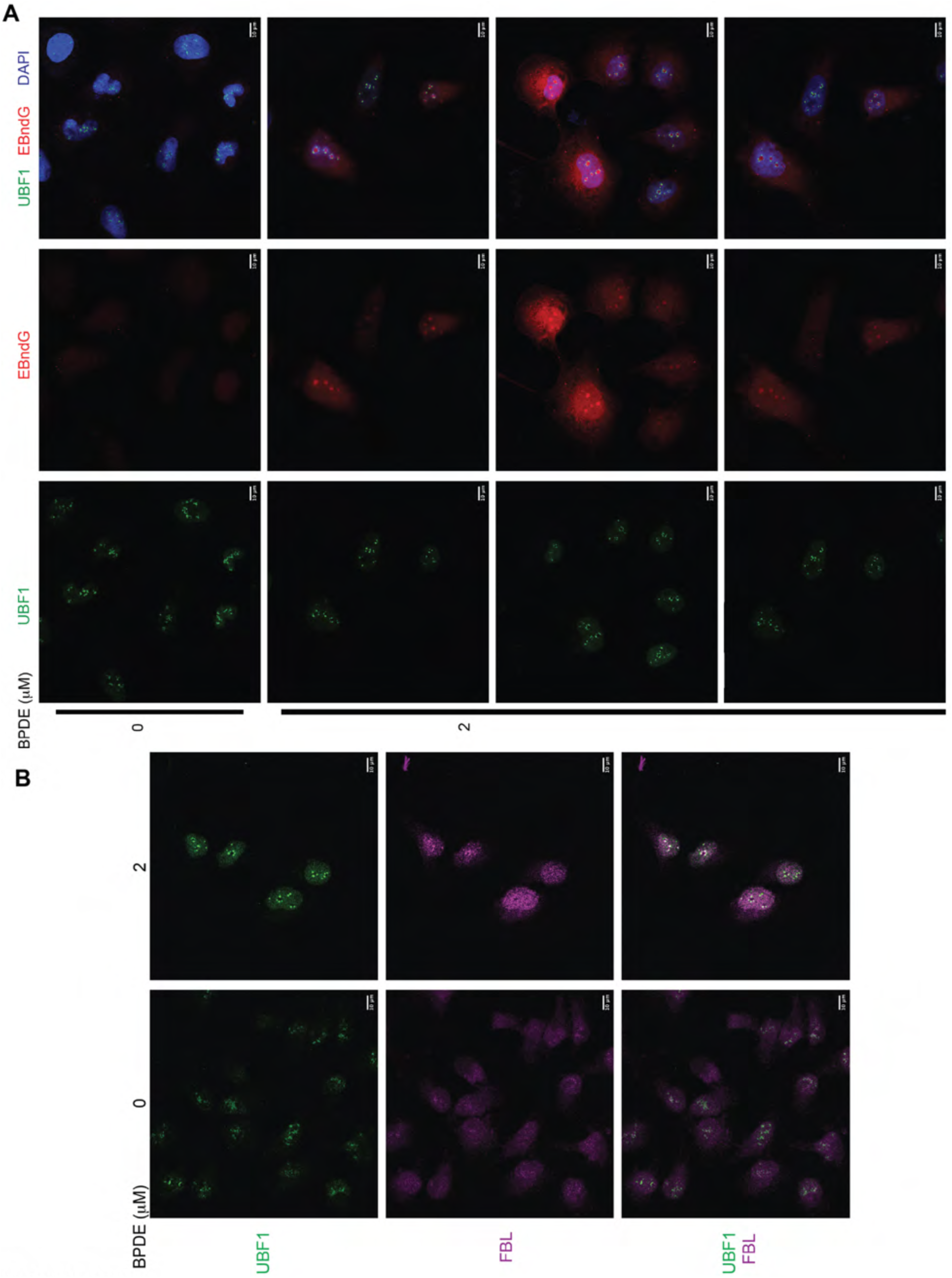
(Supplement related to Figure 2) Nucleolar stress induced by BPDE increases the activity of Polκ in the nucleolus. **A.** Full image panels of Fig 2D. **B.** Full image panels of Fig 2E. Individual UBF1 and FBL panels are shown. Scale bars = 10 μm

**Figure S4.**
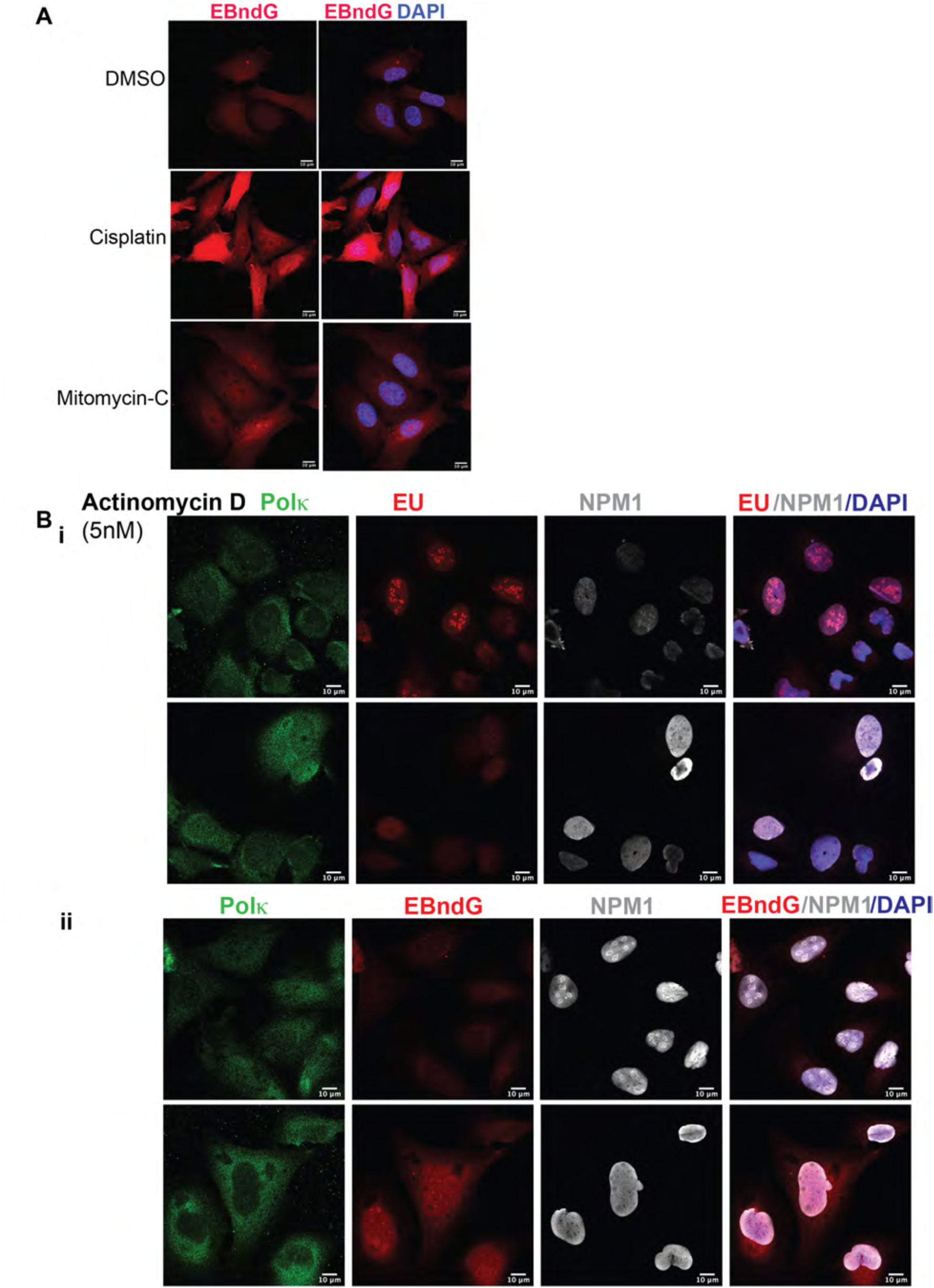
(Supplement related to Figure 3) **A.** Full image panels of Fig 3A. **B.** Full image panels of Fig 3Bi-ii.

**Figure S5.**
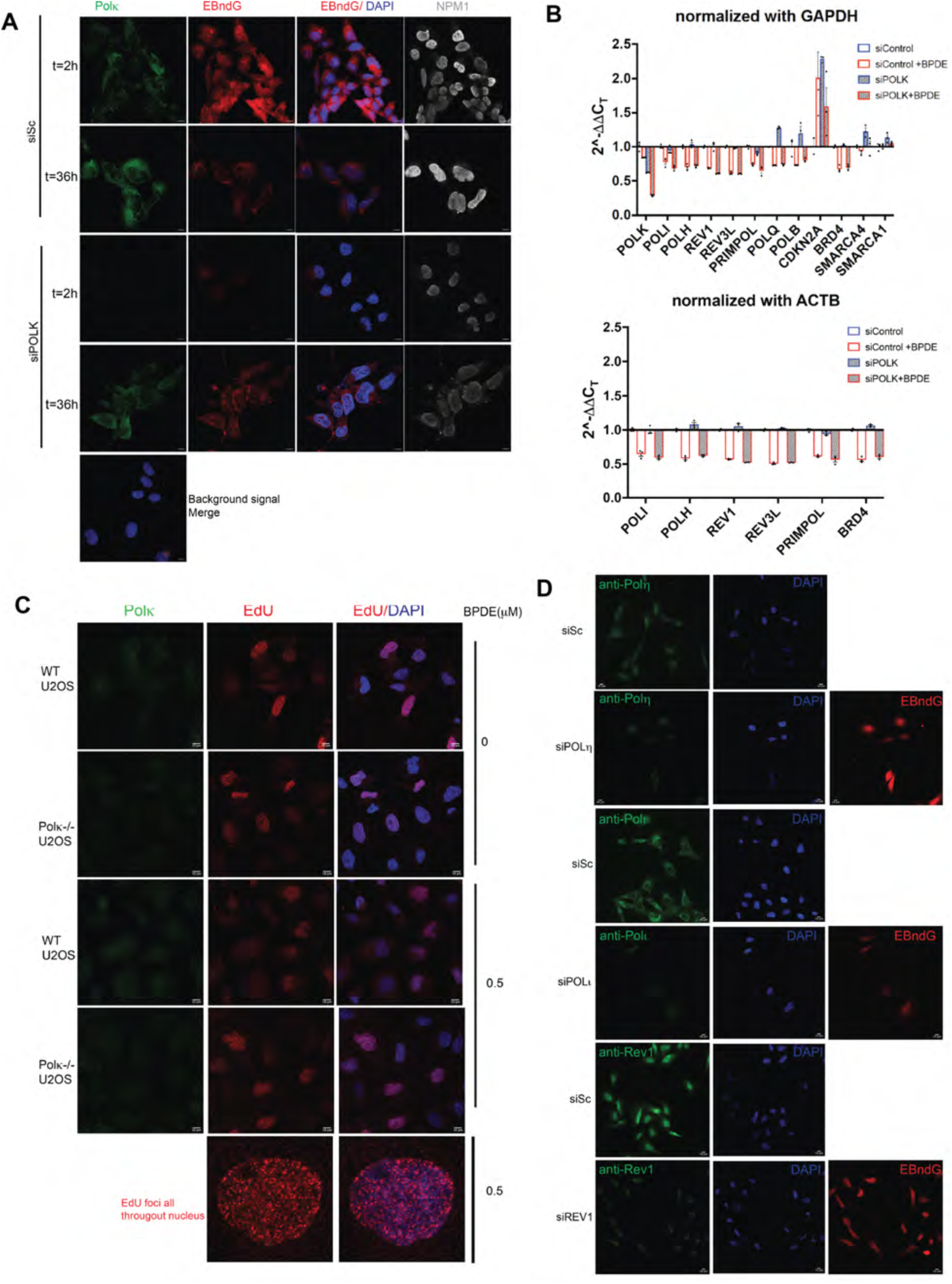
(Supplement related to Figure 4) Polκ dependent incorporation of EBndG in the nucleus and nucleolus. **A.** (Supplement for Fig 4B and 9F). Images of cells from two independent experiments transfected with siScrambled or siPOLK, treated with BPDE (0.5 μM) and fixed for immunofluorescence (t=2 h) or recovered after t=36 h. Immunofluorescence showing NPM1 (gray), Polκ (green), EBndG (red) and DAPI (blue). **B.** Relative expression of multiple transcripts after treatment with 0 and 2 μM BPDE. GAPDH or ACTB mRNA as an internal control. Calculated with the comparative CT method (mean ± SEM). All comparisons are relative to 0 μM BPDE. **C.** (Supplement for Fig 4E). EdU (in red) incorporation in WT Polκ and Polk -/- cells, after treatment with 0 and 0.5 μM BPDE, DAPI (in blue), Polκ (green). Magnified cell showing EdU foci (red). **D.** Full image panels of Fig 4F. EBndG (red) intensity compared between siScrambled RNA transfected cells from one coverslip with siPOLι, siPOLη, siREV1 transfected cells. Immunofluorescence of each knock-down cells shown in green. Scale bars = 10 μm

**Figure S6:**
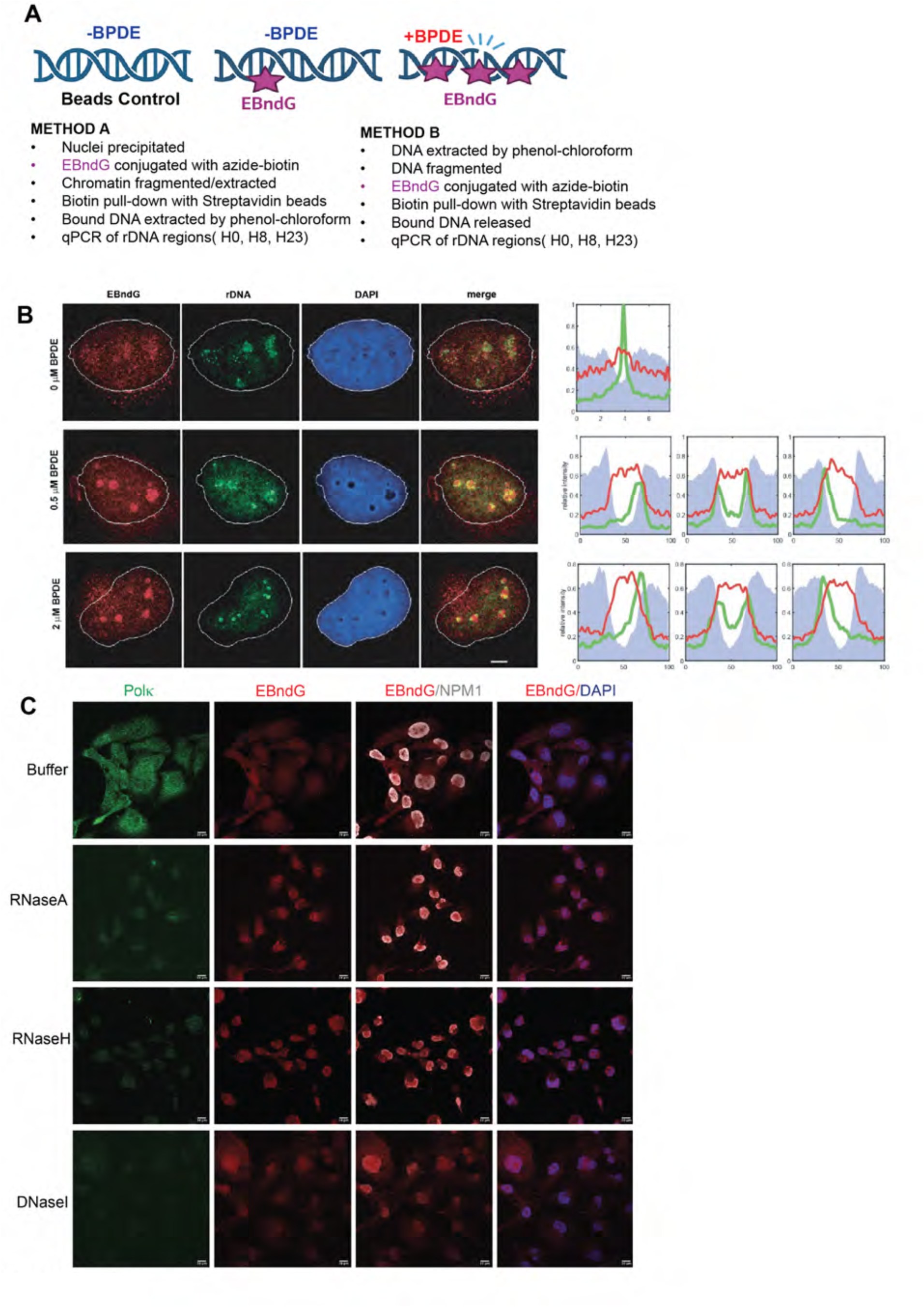
(Supplement for Figure 5). EBndG incorporation in nucleolar DNA. **A.** Schematic of the two methods used to pull down POLK-associated DNA. **B.** (Left) Representative microscopy images of cells treated with 0, 0.5 μM, 2 μM BPDE followed by 100 μM EBndG. EBndG (red) was visualized by Click conjugation with AlexaFluor 594, rDNA (green) was visualized by FISH -DNA bound to fluorescein. DNA (blue) was visualized by DAPI. The merge is the EBndG and FISH channels. (Right) The data represent profiles of EBndG (red), rDNA (green) and DAPI (gray) across the nucleolus. The 0 μM BPDE samples consisted of 7.8 μm segments with an rDNA foci centered. The BPDE treated samples were profiles across a DAPI-less region with either one or two rDNA foci. The single-rDNA foci were categorized to either the right or left side of the DAPI-less region. The profiles were normalized such that the midpoint of the downward slope of the DAPI signal was set to 33 and the midpoint of the upward slope of the DAPI was set at 67 and the total length of the profile normalized to 100. **C.** Full image panels of Fig 5C. Scale bars = 10 μm

**Figure S7:**
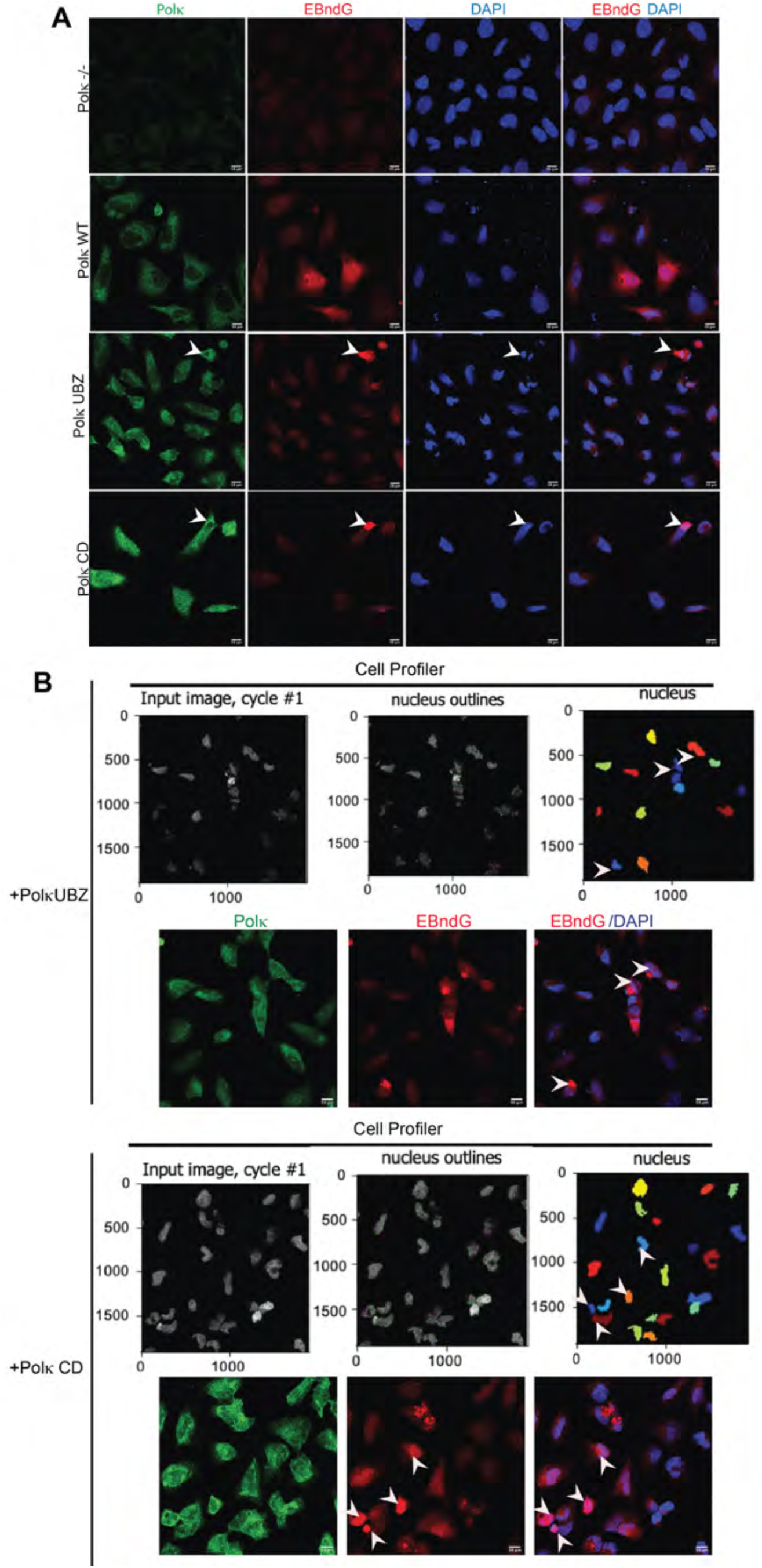
(Supplement for Figure 6B). UBZ and CD deleted Polκ with compromised activity. **A.** Full image panels of Fig 6Bi. Scale bars = 10 μm **B.** Representative image of Cell profiler pipeline, arrows showing cells with overall high fluorescence without any distinguishable nucleolar pattern, hence those cells omitted from the analysis (Fig 6Bii).

**Figure S8:**
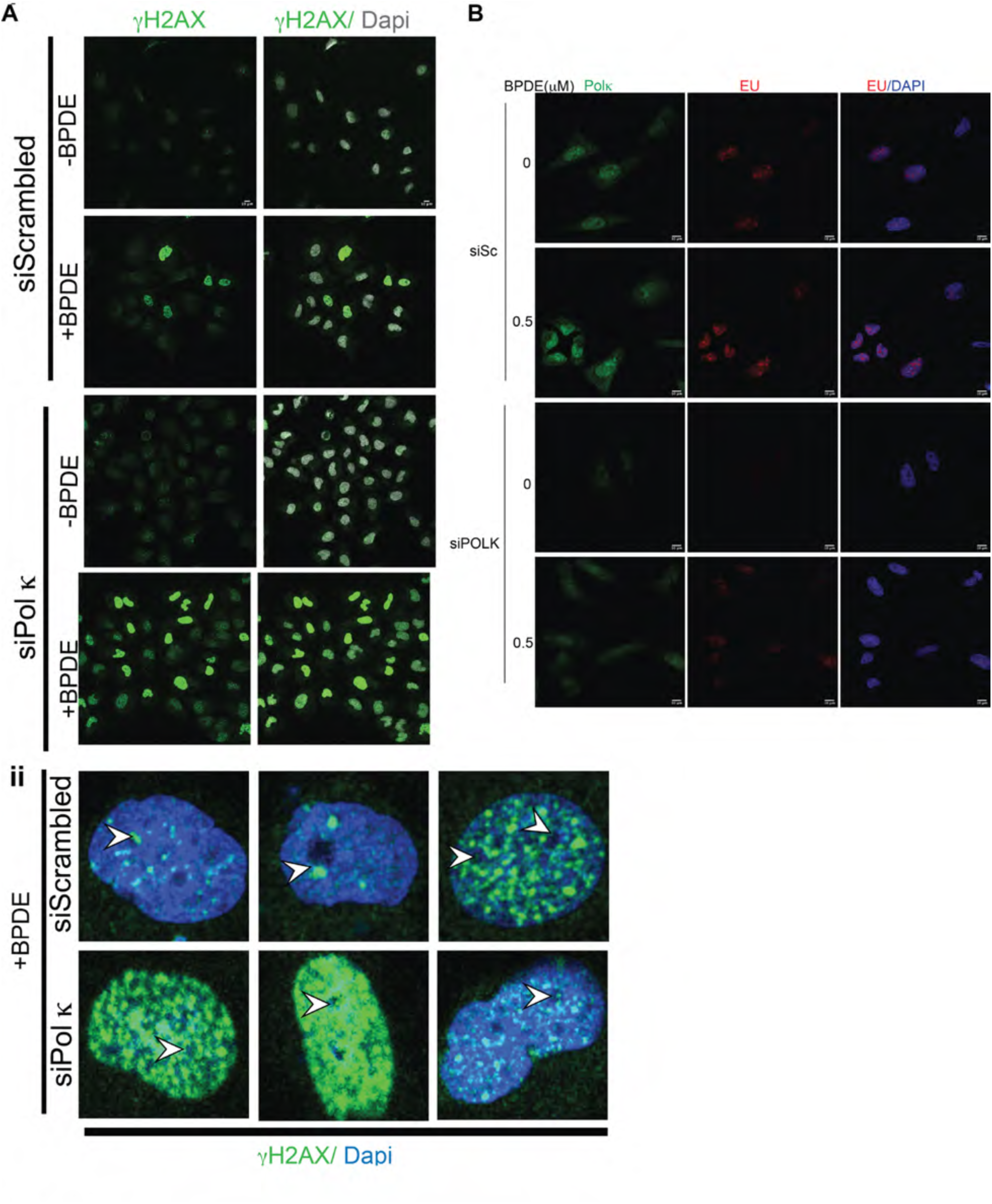
(Supplement for Figure 8) Polκ participates in recovering from nucleolar stress after BPDE damage. **A.(i)** Images for the data quantified in Figure 8A. γH2AX in green, and DAPI in gray. **(ii)** Magnified images showing γH2AX (green) in and around nucleolus pointed by arrow. **B.** Full image panels of Fig 8C. Scale bars = 10 μm

**Figure S9:**
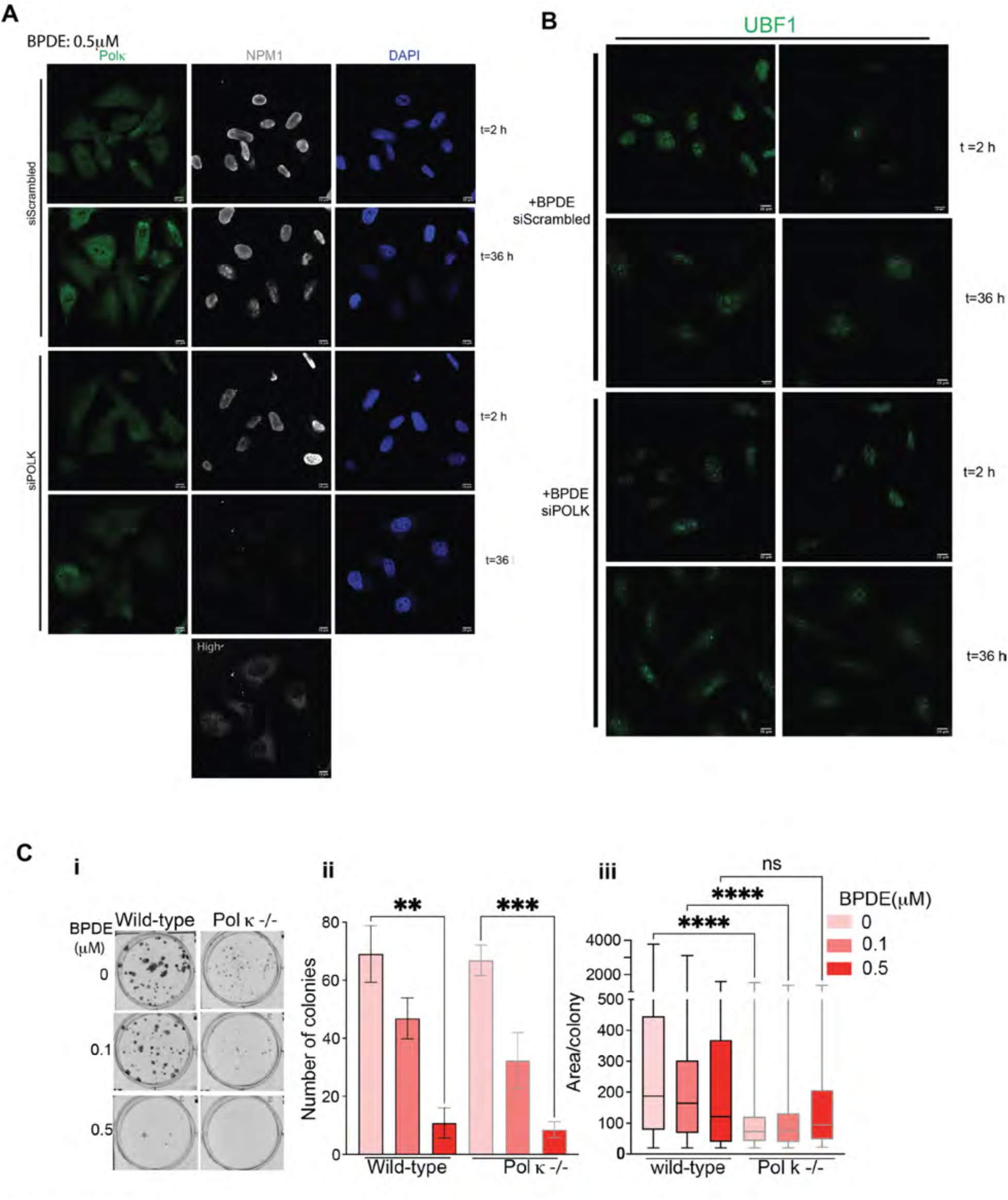
(Supplement for Figure 8) **A.** Full image panels of Fig 8F. **B.** Full image panels of Fig 8G **C.** Colony forming assays

**Figure S10:**
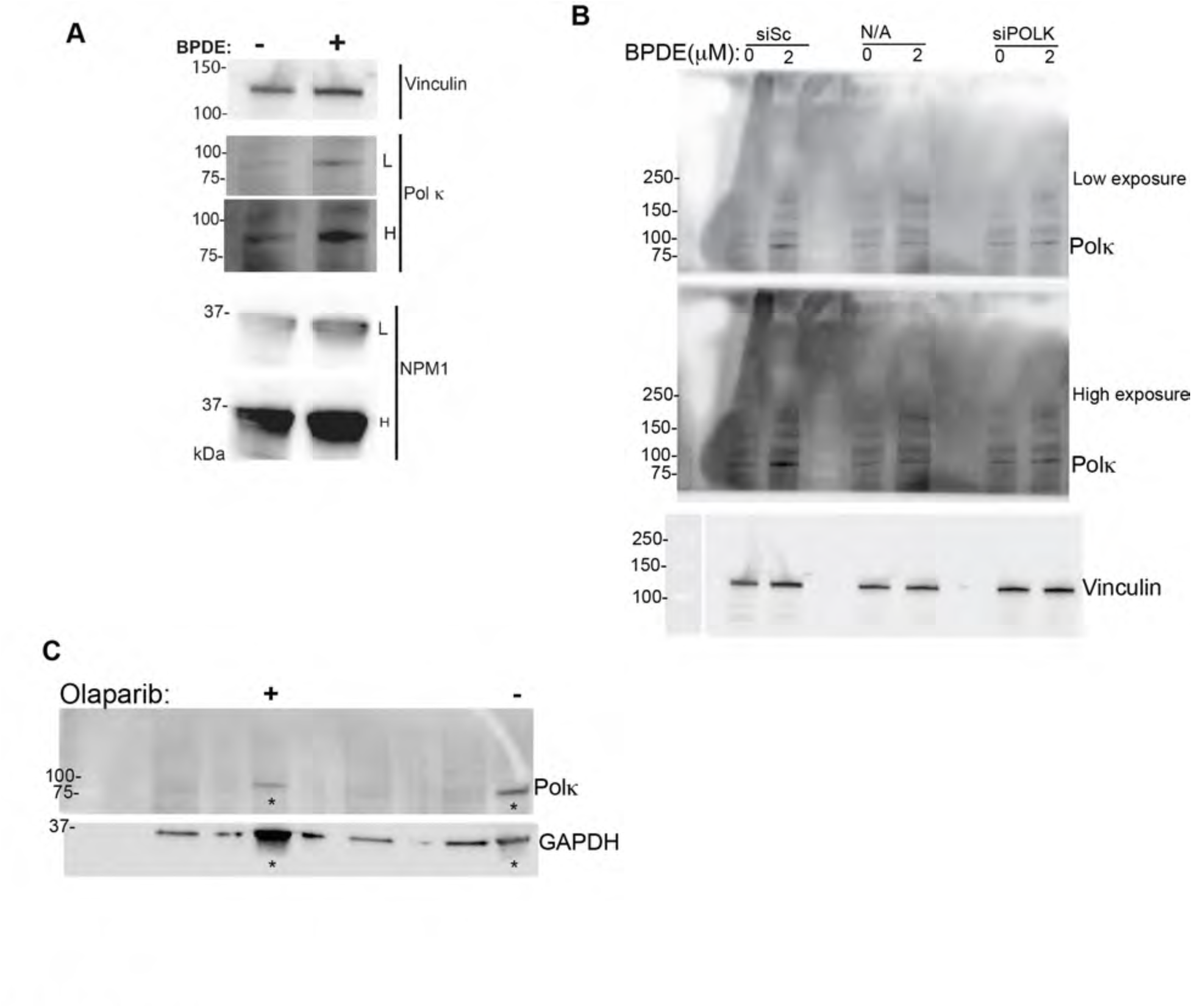
(Supplement for all western blots) **A.** Western blot analysis of Polκ and NPM1 expression (Vinculin for internal control) after treatment with 0 and 2 μM BPDE. Both low (L) and high (H) exposures are shown. **B.** Western blot analysis of Polκ expression in siScrambled (siSc) and siPOLK transfected cells. The N/A labelled lanes are irrelevant to this article. Full blot for Figure 4D **C**. Full blot for Figure 7C. The lanes with and without olaparib treatment shown in Figure 7C are marked with asterisk.

**Table S1.** (Supplement related to Figure 1) GO analysis (ToppGene) of EBndG-bound unique proteins and EdU-bound unique proteins. Biochemical Process and Chemical Component with the list of proteins.

## REFERENCES

(1) Vaisman, A.; Woodgate, R. Translesion DNA Polymerases in Eukaryotes: What Makes Them Tick? Crit Rev Biochem Mol Biol 2017, 52 (3), 274–303. 10.1080/10409238.2017.1291576.

(2) Dipple, A. DNA Adducts of Chemical Carcinogens. Carcinogenesis 1995, 16 (3), 437–441. 10.1093/carcin/16.3.437.

(3) Ohmori, H.; Ohashi, E.; Ogi, T. Mammalian Pol Kappa: Regulation of Its Expression and Lesion Substrates. Adv Protein Chem 2004, 69, 265–278. 10.1016/S0065-3233(04)69009-7.

(4) Avkin, S.; Goldsmith, M.; Velasco-Miguel, S.; Geacintov, N.; Friedberg, E. C.; Livneh, Z. Quantitative Analysis of Translesion DNA Synthesis across a Benzo[a]Pyrene-Guanine Adduct in Mammalian Cells: THE ROLE OF DNA POLYMERASE κ *. Journal of Biological Chemistry 2004, 279 (51), 53298–53305. 10.1074/jbc.M409155200.

(5) Shachar, S.; Ziv, O.; Avkin, S.; Adar, S.; Wittschieben, J.; Reißner, T.; Chaney, S.; Friedberg, E. C.; Wang, Z.; Carell, T.; Geacintov, N.; Livneh, Z. Two-Polymerase Mechanisms Dictate Error-Free and Error-Prone Translesion DNA Synthesis in Mammals. EMBO J 2009, 28 (4), 383–393. 10.1038/emboj.2008.281.

(6) Jha, V.; Ling, H. Structural Basis for Human DNA Polymerase Kappa to Bypass Cisplatin Intrastrand Cross-Link (Pt-GG) Lesion as an Efficient and Accurate Extender. J Mol Biol 2018, 430 (11), 1577–1589. 10.1016/j.jmb.2018.04.023.

(7) Stern, H. R.; Sefcikova, J.; Chaparro, V. E.; Beuning, P. J. Mammalian DNA Polymerase Kappa Activity and Specificity. 2019, 20.

(8) Bi, X.; Barkley, L. R.; Slater, D. M.; Tateishi, S.; Yamaizumi, M.; Ohmori, H.; Vaziri, C. Rad18 Regulates DNA Polymerase Kappa and Is Required for Recovery from S-Phase Checkpoint-Mediated Arrest. Mol Cell Biol 2006, 26 (9), 3527–3540. 10.1128/MCB.26.9.3527-3540.2006.

(9) Ogi, T.; Lehmann, A. R. The Y-Family DNA Polymerase κ (Pol κ) Functions in Mammalian Nucleotide-Excision Repair. Nat Cell Biol 2006, 8 (6), 640–642. 10.1038/ncb1417.

(10) Rebok, A.; Torres, M. C.; Ambrose, J. R.; Spratt, T. E. Catalytic-Dependent Role of DNA Polymerase κ in Nucleotide Excision Repair. Chem Res Toxicol 2025, 38 (6), 1103–1112. 10.1021/acs.chemrestox.5c00085.

(11) Minko, I. G.; Yamanaka, K.; Kozekov, I. D.; Kozekova, A.; Indiani, C.; O’Donnell, M. E.; Jiang, Q.; Goodman, M. F.; Rizzo, C. J.; Lloyd, R. S. Replication Bypass of the Acrolein-Mediated Deoxyguanine DNA-Peptide Cross-Links by DNA Polymerases of the DinB Family. Chem Res Toxicol 2008, 21 (10), 1983–1990. 10.1021/tx800174a.

(12) Williams, H. L.; Gottesman, M. E.; Gautier, J. Replication-Independent Repair of DNA Interstrand Crosslinks. Mol Cell 2012, 47 (1), 140–147. 10.1016/j.molcel.2012.05.001.

(13) Altshuller, M.; He, X.; MacKrell, E. J.; Wernke, K. M.; Wong, J. W. H.; Sellés-Baiget, S.; Wang, T.- Y.; Chou, T.-F.; Duxin, J. P.; Balskus, E. P.; Herzon, S. B.; Semlow, D. R. The Fanconi Anemia Pathway Repairs Colibactin-Induced DNA Interstrand Cross-Links. Molecular Biology January 30, 2024. 10.1101/2024.01.30.576698.

(14) Bétous, R.; Pillaire, M.-J.; Pierini, L.; van der Laan, S.; Recolin, B.; Ohl-Séguy, E.; Guo, C.; Niimi, N.; Grúz, P.; Nohmi, T.; Friedberg, E.; Cazaux, C.; Maiorano, D.; Hoffmann, J.-S. DNA Polymerase κ- Dependent DNA Synthesis at Stalled Replication Forks Is Important for CHK1 Activation. The EMBO Journal 2013, 32 (15), 2172–2185. 10.1038/emboj.2013.148.

(15) Baptiste, B. A.; Eckert, K. A. DNA Polymerase Kappa Microsatellite Synthesis: Two Distinct Mechanisms of Slippage-Mediated Errors. Environmental and Molecular Mutagenesis 2012, 53 (9), 787–796. 10.1002/em.21721.

(16) Walsh, E.; Wang, X.; Lee, M. Y.; Eckert, K. A. Mechanism of Replicative DNA Polymerase Delta Pausing and a Potential Role for DNA Polymerase Kappa in Common Fragile Site Replication. J Mol Biol 2013, 425 (2), 232–243. 10.1016/j.jmb.2012.11.016.

(17) Tang, F.; Wang, Y.; Zhao, T.; Yuan, J.; Kellum, A. H.; Wang, Y. DNA Polymerase κ Participates in Early S-Phase DNA Replication in Human Cells. Proc Natl Acad Sci U S A 2024, 121 (28), e2405473121. 10.1073/pnas.2405473121.

(18) Torres, M. C.; Rebok, A.; Sun, D.; Spratt, T. E. Activity of DNA Polymerase κ across the Genome in Human Fibroblasts. Proc Natl Acad Sci U S A 2024, 121 (28), e2403130121. 10.1073/pnas.2403130121.

(19) Prakasha Gowda, A. S.; Lee, M.; Spratt, T. E. N2-Substituted 2ʹ-Deoxyguanosine Triphosphate Derivatives, Selective Substrates for Human DNA Polymerase κ. Angew Chem Int Ed Engl 2017, 56 (10), 2628–2631. 10.1002/anie.201611607.

(20) Henderson, A. S.; Warburton, D.; Atwood, K. C. Location of Ribosomal DNA in the Human Chromosome Complement. Proc Natl Acad Sci U S A 1972, 69 (11), 3394–3398. 10.1073/pnas.69.11.3394.

(21) van Sluis, M.; van Vuuren, C.; Mangan, H.; McStay, B. NORs on Human Acrocentric Chromosome P-Arms Are Active by Default and Can Associate with Nucleoli Independently of rDNA. Proc Natl Acad Sci U S A 2020, 117 (19), 10368–10377. 10.1073/pnas.2001812117.

(22) Warner, J. R. The Economics of Ribosome Biosynthesis in Yeast. Trends Biochem Sci 1999, 24 (11), 437–440. 10.1016/s0968-0004(99)01460-7.

(23) Lewis, J. D.; Tollervey, D. Like Attracts like: Getting RNA Processing Together in the Nucleus. Science 2000, 288 (5470), 1385–1389. 10.1126/science.288.5470.1385.

(24) McStay, B. Nucleolar Organizer Regions: Genomic “dark Matter” Requiring Illumination. Genes Dev 2016, 30 (14), 1598–1610. 10.1101/gad.283838.116.

(25) Rudra, D.; Warner, J. R. What Better Measure than Ribosome Synthesis? Genes Dev. 2004, 18 (20), 2431–2436. 10.1101/gad.1256704.

(26) Cerqueira, A. V.; Lemos, B. Ribosomal DNA and the Nucleolus as Keystones in Nuclear Architecture, Organization, and Function. Trends Genet 2019, 35 (10), 710–723. 10.1016/j.tig.2019.07.011.

(27) Pederson, T. The Nucleolus. Cold Spring Harb Perspect Biol 2011, 3 (3), a000638. 10.1101/cshperspect.a000638.

(28) Farley, K. I.; Surovtseva, Y.; Merkel, J.; Baserga, S. J. Determinants of Mammalian Nucleolar Architecture. Chromosoma 2015, 124 (3), 323–331. 10.1007/s00412-015-0507-z.

(29) Aguilera, A.; Gómez-González, B. Genome Instability: A Mechanistic View of Its Causes and Consequences. Nat Rev Genet 2008, 9 (3), 204–217. 10.1038/nrg2268.

(30) Li, X.; Manley, J. L. Cotranscriptional Processes and Their Influence on Genome Stability. Genes Dev. 2006, 20 (14), 1838–1847. 10.1101/gad.1438306.

(31) Gan, W.; Guan, Z.; Liu, J.; Gui, T.; Shen, K.; Manley, J. L.; Li, X. R-Loop-Mediated Genomic Instability Is Caused by Impairment of Replication Fork Progression. Genes Dev. 2011, 25 (19), 2041– 2056. 10.1101/gad.17010011.

(32) Lindström, M. S.; Jurada, D.; Bursac, S.; Orsolic, I.; Bartek, J.; Volarevic, S. Nucleolus as an Emerging Hub in Maintenance of Genome Stability and Cancer Pathogenesis. Oncogene 2018, 37 (18), 2351–2366. 10.1038/s41388-017-0121-z.

(33) Kruhlak, M.; Crouch, E. E.; Orlov, M.; Montaño, C.; Gorski, S. A.; Nussenzweig, A.; Misteli, T.; Phair, R. D.; Casellas, R. The ATM Repair Pathway Inhibits RNA Polymerase I Transcription in Response to Chromosome Breaks. Nature 2007, 447 (7145), 730–734. 10.1038/nature05842.

(34) van Sluis, M.; McStay, B. A Localized Nucleolar DNA Damage Response Facilitates Recruitment of the Homology-Directed Repair Machinery Independent of Cell Cycle Stage. Genes Dev 2015, 29 (11), 1151–1163. 10.1101/gad.260703.115.

(35) van Sluis, M.; McStay, B. Nucleolar Reorganization in Response to rDNA Damage. Curr Opin Cell Biol 2017, 46, 81–86. 10.1016/j.ceb.2017.03.004.

(36) van Sluis, M.; McStay, B. Nucleolar DNA Double-Strand Break Responses Underpinning rDNA Genomic Stability. Trends Genet 2019, 35 (10), 743–753. 10.1016/j.tig.2019.07.001.

(37) Boulon, S.; Westman, B. J.; Hutten, S.; Boisvert, F.-M.; Lamond, A. I. The Nucleolus under Stress. Mol Cell 2010, 40 (2), 216–227. 10.1016/j.molcel.2010.09.024.

(38) Lindström, M. S. NPM1/B23: A Multifunctional Chaperone in Ribosome Biogenesis and Chromatin Remodeling. Biochem Res Int 2011, 2011, 195209. 10.1155/2011/195209.

(39) Rancourt, A.; Satoh, M. S. Delocalization of Nucleolar Poly(ADP-Ribose) Polymerase-1 to the Nucleoplasm and Its Novel Link to Cellular Sensitivity to DNA Damage. DNA Repair (Amst) 2009, 8 (3), 286–297. 10.1016/j.dnarep.2008.11.018.

(40) Yung, T. M. C.; Sato, S.; Satoh, M. S. Poly(ADP-Ribosyl)Ation as a DNA Damage-Induced Post-Translational Modification Regulating Poly(ADP-Ribose) Polymerase-1-Topoisomerase I Interaction. J Biol Chem 2004, 279 (38), 39686–39696. 10.1074/jbc.M402729200.

(41) Yang, Y.; Hu, J.; Selby, C. P.; Li, W.; Yimit, A.; Jiang, Y.; Sancar, A. Single-Nucleotide Resolution Analysis of Nucleotide Excision Repair of Ribosomal DNA in Humans and Mice. Journal of Biological Chemistry 2019, 294 (1), 210–217. 10.1074/jbc.RA118.006121.

(42) Daniel, L.; Cerutti, E.; Donnio, L.-M.; Nonnekens, J.; Carrat, C.; Zahova, S.; Mari, P.-O.; Giglia-Mari, G. Mechanistic Insights in Transcription-Coupled Nucleotide Excision Repair of Ribosomal DNA. Proceedings of the National Academy of Sciences 2018, 115 (29), E6770–E6779. 10.1073/pnas.1716581115.

(43) Ahmad, Y.; Boisvert, F.-M.; Gregor, P.; Cobley, A.; Lamond, A. I. NOPdb: Nucleolar Proteome Database--2008 Update. Nucleic Acids Res 2009, 37 (Database issue), D181–184. 10.1093/nar/gkn804.

(44) Ogawa, L. M.; Baserga, S. J. Crosstalk between the Nucleolus and the DNA Damage Response. Mol Biosyst 2017, 13 (3), 443–455. 10.1039/c6mb00740f.

(45) Moore, H. M.; Bai, B.; Boisvert, F.-M.; Latonen, L.; Rantanen, V.; Simpson, J. C.; Pepperkok, R.; Lamond, A. I.; Laiho, M. Quantitative Proteomics and Dynamic Imaging of the Nucleolus Reveal Distinct Responses to UV and Ionizing Radiation. Mol Cell Proteomics 2011, 10 (10), M111.009241. 10.1074/mcp.M111.009241.

(46) Andersen, J. S.; Lam, Y. W.; Leung, A. K. L.; Ong, S.-E.; Lyon, C. E.; Lamond, A. I.; Mann, M. Nucleolar Proteome Dynamics. Nature 2005, 433 (7021), 77–83. 10.1038/nature03207.

(47) Wang, H.; Chen, T.; Weng, C.; Yang, C.; Tang, M. Acrolein Preferentially Damages Nucleolus Eliciting Ribosomal Stress and Apoptosis in Human Cancer Cells. Oncotarget 2016, 7 (49), 80450– 80464. 10.18632/oncotarget.12608.

(48) Sutton, E. C.; DeRose, V. J. Early Nucleolar Responses Differentiate Mechanisms of Cell Death Induced by Oxaliplatin and Cisplatin. J Biol Chem 2021, 296, 100633. 10.1016/j.jbc.2021.100633.

(49) Sutton, E. C.; McDevitt, C. E.; Prochnau, J. Y.; Yglesias, M. V.; Mroz, A. M.; Yang, M. C.; Cunningham, R. M.; Hendon, C. H.; DeRose, V. J. Nucleolar Stress Induction by Oxaliplatin and Derivatives. J. Am. Chem. Soc. 2019, 141 (46), 18411–18415. 10.1021/jacs.9b10319.

(50) Thakar, T.; Leung, W.; Nicolae, C. M.; Clements, K. E.; Shen, B.; Bielinsky, A.-K.; Moldovan, G.-L. Ubiquitinated-PCNA Protects Replication Forks from DNA2-Mediated Degradation by Regulating Okazaki Fragment Maturation and Chromatin Assembly. Nat Commun 2020, 11 (1), 2147. 10.1038/s41467-020-16096-w.

(51) Tonzi, P.; Yin, Y.; Lee, C. W. T.; Rothenberg, E.; Huang, T. T. Translesion Polymerase Kappa- Dependent DNA Synthesis Underlies Replication Fork Recovery. eLife 7, e41426. 10.7554/eLife.41426.

(52) Wiest, N. E.; Tomkinson, A. E. Chapter One - Optimization of Native and Formaldehyde iPOND Techniques for Use in Suspension Cells. In Methods in Enzymology; Eichman, B. F., Ed.; DNA Repair Enzymes: Cell, Molecular, and Chemical Biology; Academic Press, 2017; Vol. 591, pp 1–32. 10.1016/bs.mie.2017.03.001.

(53) Bryant, C. J.; McCool, M. A.; Abriola, L.; Surovtseva, Y. V.; Baserga, S. J. A High-Throughput Assay for Directly Monitoring Nucleolar rRNA Biogenesis. Open Biol 2022, 12 (1), 210305. 10.1098/rsob.210305.

(54) O’Sullivan, A. C.; Sullivan, G. J.; McStay, B. UBF Binding In Vivo Is Not Restricted to Regulatory Sequences within the Vertebrate Ribosomal DNA Repeat. Mol Cell Biol 2002, 22 (2), 657–668. 10.1128/MCB.22.2.657-668.2002.

(55) van Sluis, M.; van Vuuren, C.; McStay, B. The Relationship Between Human Nucleolar Organizer Regions and Nucleoli, Probed by 3D-ImmunoFISH. In The Nucleolus: Methods and Protocols; Németh, A., Ed.; Springer: New York, NY, 2016; pp 3–14. 10.1007/978-1-4939-3792-9_1.

(56) Carpenter, A. E.; Jones, T. R.; Lamprecht, M. R.; Clarke, C.; Kang, I. H.; Friman, O.; Guertin, D. A.; Chang, J. H.; Lindquist, R. A.; Moffat, J.; Golland, P.; Sabatini, D. M. CellProfiler: Image Analysis Software for Identifying and Quantifying Cell Phenotypes. Genome Biology 2006, 7 (10), R100. 10.1186/gb-2006-7-10-r100.

(57) Leung, K. H. T.; Abou El Hassan, M.; Bremner, R. A Rapid and Efficient Method to Purify Proteins at Replication Forks under Native Conditions. Biotechniques 2013, 55 (4), 204–206. 10.2144/000114089.

(58) Nwagbara, O.; Darling-Reed, S. F.; Tucker, A.; Harris, C.; Abazinge, M.; Thomas, R. D.; Gragg, R. D. Induction of Cell Death, DNA Strand Breaks, and Cell Cycle Arrest in DU145 Human Prostate Carcinoma Cell Line by Benzo[a]Pyrene and Benzo[a]Pyrene-7,8-Diol-9,10-Epoxide. Int J Environ Res Public Health 2007, 4 (1), 10–14. 10.3390/ijerph2007010002.

(59) Dreij, K.; Rhrissorrakrai, K.; Gunsalus, K. C.; Geacintov, N. E.; Scicchitano, D. A. Benzo[a]Pyrene Diol Epoxide Stimulates an Inflammatory Response in Normal Human Lung Fibroblasts through a P53 and JNK Mediated Pathway. Carcinogenesis 2010, 31 (6), 1149–1157. 10.1093/carcin/bgq073.

(60) Fischer, J. M. F.; Zubel, T.; Jander, K.; Fix, J.; Trussina, I. R. E. A.; Gebhard, D.; Bergemann, J.; Bürkle, A.; Mangerich, A. PARP1 Protects from Benzo[a]Pyrene Diol Epoxide-Induced Replication Stress and Mutagenicity. Arch Toxicol 2018, 92 (3), 1323–1340. 10.1007/s00204-017-2115-6.

(61) Alabert, C.; Bukowski-Wills, J. C.; Lee, S. B.; Kustatscher, G.; Nakamura, K.; De Lima Alves, F.; Menard, P.; Mejlvang, J.; Rappsilber, J.; Groth, A. Nascent Chromatin Capture Proteomics Determines Chromatin Dynamics during DNA Replication and Identifies Unknown Fork Components. Nature Cell Biology 2014, 16 (3), 281–291. 10.1038/ncb2918.

(62) Dungrawala, H.; Rose, K. L.; Bhat, K. P.; Mohni, K. N.; Glick, G. G.; Couch, F. B.; Cortez, D. The Replication Checkpoint Prevents Two Types of Fork Collapse without Regulating Replisome Stability. Mol Cell 2015, 59 (6), 998–1010. 10.1016/j.molcel.2015.07.030.

(63) Wessel, S. R.; Mohni, K. N.; Luzwick, J. W.; Dungrawala, H.; Cortez, D. Functional Analysis of the Replication Fork Proteome Identifies BET Proteins as PCNA Regulators. Cell Rep 2019, 28 (13), 3497–3509.e4. 10.1016/j.celrep.2019.08.051.

(64) Romagnolo, D.; Jeffy, B.; Samuelson, D.; Payne, C. Microarray Expression Profiles in Breast Cancer MCF-7 Cells Following Exposure to Benzo[a]Pyrene and Its Diol Epoxide. Nat Genet 2001, 27 (4), 83–83. 10.1038/87272.

(65) Zhuo, M.; Gorgun, M. F.; Englander, E. W. Translesion Synthesis DNA Polymerase Kappa Is Indispensable for DNA Repair Synthesis in Cisplatin Exposed Dorsal Root Ganglion Neurons. Mol Neurobiol 2018, 55 (3), 2506–2515. 10.1007/s12035-017-0507-5.

(66) Spanjaard, A.; Shah, R.; de Groot, D.; Buoninfante, O. A.; Morris, B.; Lieftink, C.; Pritchard, C.; Zürcher, L. M.; Ormel, S.; Catsman, J. J. I.; de Korte-Grimmerink, R.; Siteur, B.; Proost, N.; Boadum, T.; van de Ven, M.; Song, J.-Y.; Kreft, M.; van den Berk, P. C. M.; Beijersbergen, R. L.; Jacobs, H. Division of Labor within the DNA Damage Tolerance System Reveals Non-Epistatic and Clinically Actionable Targets for Precision Cancer Medicine. Nucleic Acids Research 2022, 50 (13), 7420–7435. 10.1093/nar/gkac545.

(67) Kanemaru, Y.; Suzuki, T.; Sassa, A.; Matsumoto, K.; Adachi, N.; Honma, M.; Numazawa, S.; Nohmi, T. DNA Polymerase Kappa Protects Human Cells against MMC-Induced Genotoxicity through Error-Free Translesion DNA Synthesis. Genes Environ 2017, 39, 6. 10.1186/s41021-016-0067-3.

(68) Takeiri, A.; Wada, N. A.; Motoyama, S.; Matsuzaki, K.; Tateishi, H.; Matsumoto, K.; Niimi, N.; Sassa, A.; Grúz, P.; Masumura, K.; Yamada, M.; Mishima, M.; Jishage, K.; Nohmi, T. In Vivo Evidence That DNA Polymerase Kappa Is Responsible for Error-Free Bypass across DNA Cross-Links Induced by Mitomycin C. DNA Repair 2014, 24, 113–121. 10.1016/j.dnarep.2014.09.002.

(69) Harding, S. M.; Boiarsky, J. A.; Greenberg, R. A. ATM Dependent Silencing Links Nucleolar Chromatin Reorganization to DNA Damage Recognition. Cell Rep 2015, 13 (2), 251–259. 10.1016/j.celrep.2015.08.085.

(70) Schoefl, G. I. The Effect of Actinomycin D on the Fine Structure of the Nucleolus. Journal of Ultrastructure Research 1964, 10 (3), 224–243. 10.1016/S0022-5320(64)80007-1.

(71) Peltonen, K.; Colis, L.; Liu, H.; Trivedi, R.; Moubarek, M. S.; Moore, H. M.; Bai, B.; Rudek, M. A.; Bieberich, C. J.; Laiho, M. A Targeting Modality for Destruction of RNA Polymerase I That Possesses Anticancer Activity. Cancer Cell 2014, 25 (1), 77–90. 10.1016/j.ccr.2013.12.009.

(72) Schöfer, C.; Weipoltshammer, K.; Almeder, M.; Müller, M.; Wachtler, F. Redistribution of Ribosomal DNA after Blocking of Transcription Induced by Actinomycin D. Chromosome Res 1996, 4 (5), 384–391. 10.1007/BF02257274.

(73) Reynolds, R. C.; Montgomery, P. O.; Hughes, B. NUCLEOLAR “CAPS” PRODUCED BY ACTINOMYCIN D. Cancer Res 1964, 24, 1269–1277.

(74) Tornaletti, S. Transcription Arrest at DNA Damage Sites. Mutation Research/Fundamental and Molecular Mechanisms of Mutagenesis 2005, 577 (1), 131–145. 10.1016/j.mrfmmm.2005.03.014.

(75) Geijer, M. E.; Marteijn, J. A. What Happens at the Lesion Does Not Stay at the Lesion: Transcription-Coupled Nucleotide Excision Repair and the Effects of DNA Damage on Transcription in Cis and Trans. DNA Repair (Amst) 2018, 71, 56–68. 10.1016/j.dnarep.2018.08.007.

(76) Steurer, B.; Janssens, R. C.; Geijer, M. E.; Aprile-Garcia, F.; Geverts, B.; Theil, A. F.; Hummel, B.; van Royen, M. E.; Evers, B.; Bernards, R.; Houtsmuller, A. B.; Sawarkar, R.; Marteijn, J. DNA Damage-Induced Transcription Stress Triggers the Genome-Wide Degradation of Promoter-Bound Pol II. Nat Commun 2022, 13 (1), 3624. 10.1038/s41467-022-31329-w.

(77) Brueckner, F.; Hennecke, U.; Carell, T.; Cramer, P. CPD Damage Recognition by Transcribing RNA Polymerase II. Science 2007, 315 (5813), 859–862. 10.1126/science.1135400.

(78) Ziv, O.; Zeisel, A.; Mirlas-Neisberg, N.; Swain, U.; Nevo, R.; Ben-Chetrit, N.; Martelli, M. P.; Rossi, R.; Schiesser, S.; Canman, C. E.; Carell, T.; Geacintov, N. E.; Falini, B.; Domany, E.; Livneh, Z. Identification of Novel DNA-Damage Tolerance Genes Reveals Regulation of Translesion DNA Synthesis by Nucleophosmin. Nat Commun 2014, 5 (1), 5437. 10.1038/ncomms6437.

(79) Dhoonmoon, A.; Ambrose, J. R.; Garg, S.; Lascarez-Espana, C.; Rebok, A.; Spratt, T. E.; Moldovan, G.-L.; Nicolae, C. M. Translesion-Synthesis-Mediated Bypass of DNA Lesions Occurs Predominantly behind Replication Forks Restarted by PrimPol. Cell Rep 2025, 44 (3), 115360. 10.1016/j.celrep.2025.115360.

(80) Bi, X.; Slater, D. M.; Ohmori, H.; Vaziri, C. DNA Polymerase Kappa Is Specifically Required for Recovery from the Benzo[a]Pyrene-Dihydrodiol Epoxide (BPDE)-Induced S-Phase Checkpoint. J Biol Chem 2005, 280 (23), 22343–22355. 10.1074/jbc.M501562200.

(81) Fan, W.; Eklund, E.; Sherman, R. M.; Liu, H.; Pitts, S.; Ford, B.; Rajeshkumar, N. V.; Laiho, M. Widespread Genetic Heterogeneity of Human Ribosomal RNA Genes. RNA 2022, 28 (4), 478–492. 10.1261/rna.078925.121.

(82) Chaffer, C. L.; Brueckmann, I.; Scheel, C.; Kaestli, A. J.; Wiggins, P. A.; Rodrigues, L. O.; Brooks, M.; Reinhardt, F.; Su, Y.; Polyak, K.; Arendt, L. M.; Kuperwasser, C.; Bierie, B.; Weinberg, R. A. Normal and Neoplastic Nonstem Cells Can Spontaneously Convert to a Stem-like State. Proc Natl Acad Sci U S A 2011, 108 (19), 7950–7955. 10.1073/pnas.1102454108.

(83) Zentner, G. E.; Saiakhova, A.; Manaenkov, P.; Adams, M. D.; Scacheri, P. C. Integrative Genomic Analysis of Human Ribosomal DNA. Nucleic Acids Res 2011, 39 (12), 4949–4960. 10.1093/nar/gkq1326.

(84) Németh, A.; Guibert, S.; Tiwari, V. K.; Ohlsson, R.; Längst, G. Epigenetic Regulation of TTF-I-mediated Promoter–Terminator Interactions of rRNA Genes. EMBO J 2008, 27 (8), 1255–1265. 10.1038/emboj.2008.57.

(85) Dimitrova, D. S. DNA Replication Initiation Patterns and Spatial Dynamics of the Human Ribosomal RNA Gene Loci. J Cell Sci 2011, 124 (16), 2743–2752. 10.1242/jcs.082230.

(86) Schlesinger, S.; Selig, S.; Bergman, Y.; Cedar, H. Allelic Inactivation of rDNA Loci. Genes Dev. 2009, 23 (20), 2437–2447. 10.1101/gad.544509.

(87) Warmerdam, D. O.; Wolthuis, R. M. F. Keeping Ribosomal DNA Intact: A Repeating Challenge. Chromosome Res 2019, 27 (1), 57–72. 10.1007/s10577-018-9594-z.

(88) Horigome, C.; Unozawa, E.; Ooki, T.; Kobayashi, T. Ribosomal RNA Gene Repeats Associate with the Nuclear Pore Complex for Maintenance after DNA Damage. PLoS Genet 2019, 15 (4), e1008103. 10.1371/journal.pgen.1008103.

(89) Salim, D.; Gerton, J. L. Ribosomal DNA Instability and Genome Adaptability. Chromosome Res 2019, 27 (1–2), 73–87. 10.1007/s10577-018-9599-7.

(90) Murayama, A.; Ohmori, K.; Fujimura, A.; Minami, H.; Yasuzawa-Tanaka, K.; Kuroda, T.; Oie, S.; Daitoku, H.; Okuwaki, M.; Nagata, K.; Fukamizu, A.; Kimura, K.; Shimizu, T.; Yanagisawa, J. Epigenetic Control of rDNA Loci in Response to Intracellular Energy Status. Cell 2008, 133 (4), 627–639. 10.1016/j.cell.2008.03.030.

(91) Hoege, C.; Pfander, B.; Moldovan, G.-L.; Pyrowolakis, G.; Jentsch, S. RAD6-Dependent DNA Repair Is Linked to Modification of PCNA by Ubiquitin and SUMO. Nature 2002, 419 (6903), 135–141. 10.1038/nature00991.

(92) Lehmann, A. R.; Niimi, A.; Ogi, T.; Brown, S.; Sabbioneda, S.; Wing, J. F.; Kannouche, P. L.; Green, C. M. Translesion Synthesis: Y-Family Polymerases and the Polymerase Switch. DNA Repair (Amst) 2007, 6 (7), 891–899. 10.1016/j.dnarep.2007.02.003.

(93) Ghosal, G.; Chen, J. DNA Damage Tolerance: A Double-Edged Sword Guarding the Genome. Translational Cancer Research 2013, 2 (3). 10.3978/j.issn.2218-676X.2013.04.01.

(94) Choe, K. N.; Moldovan, G.-L. Forging Ahead through Darkness: PCNA, Still the Principal Conductor at the Replication Fork. Mol Cell 2017, 65 (3), 380–392. 10.1016/j.molcel.2016.12.020.

(95) Jones, M. J.; Colnaghi, L.; Huang, T. T. Dysregulation of DNA Polymerase κ Recruitment to Replication Forks Results in Genomic Instability. The EMBO Journal 2012, 31 (4), 908–918. 10.1038/emboj.2011.457.

(96) Kannouche, P. L.; Wing, J.; Lehmann, A. R. Interaction of Human DNA Polymerase Eta with Monoubiquitinated PCNA: A Possible Mechanism for the Polymerase Switch in Response to DNA Damage. Mol Cell 2004, 14 (4), 491–500. 10.1016/s1097-2765(04)00259-x.

(97) Bienko, M.; Green, C. M.; Crosetto, N.; Rudolf, F.; Zapart, G.; Coull, B.; Kannouche, P.; Wider, G.; Peter, M.; Lehmann, A. R.; Hofmann, K.; Dikic, I. Ubiquitin-Binding Domains in Y-Family Polymerases Regulate Translesion Synthesis. Science 2005, 310 (5755), 1821–1824. 10.1126/science.1120615.

(98) Ohmori, H.; Hanafusa, T.; Ohashi, E.; Vaziri, C. Separate Roles of Structured and Unstructured Regions of Y-Family DNA Polymerases. In Advances in Protein Chemistry and Structural Biology; McPherson, A., Ed.; Advances in Protein Chemistry and Structural Biology; Academic Press, 2009; Vol. 78, pp 99–146. 10.1016/S1876-1623(08)78004-0.

(99) Gerlach, V. L.; Feaver, W. J.; Fischhaber, P. L.; Friedberg, E. C. Purification and Characterization of Polκ, a DNA Polymerase Encoded by the Human DINB1 Gene *. Journal of Biological Chemistry 2001, 276 (1), 92–98. 10.1074/jbc.M004413200.

(100) Lancey, C.; Tehseen, M.; Bakshi, S.; Percival, M.; Takahashi, M.; Sobhy, M. A.; Raducanu, V. S.; Blair, K.; Muskett, F. W.; Ragan, T. J.; Crehuet, R.; Hamdan, S. M.; De Biasio, A. Cryo-EM Structure of Human Pol κ Bound to DNA and Mono-Ubiquitylated PCNA. Nat Commun 2021, 12 (1), 6095. 10.1038/s41467-021-26251-6.

(101) Lone, S.; Townson, S. A.; Uljon, S. N.; Johnson, R. E.; Brahma, A.; Nair, D. T.; Prakash, S.; Prakash, L.; Aggarwal, A. K. Human DNA Polymerase κ Encircles DNA: Implications for Mismatch Extension and Lesion Bypass. Molecular Cell 2007, 25 (4), 601–614. 10.1016/j.molcel.2007.01.018.

(102) Gagné, J.-P.; Isabelle, M.; Lo, K. S.; Bourassa, S.; Hendzel, M. J.; Dawson, V. L.; Dawson, T. M.; Poirier, G. G. Proteome-Wide Identification of Poly(ADP-Ribose) Binding Proteins and Poly(ADP-Ribose)-Associated Protein Complexes. Nucleic Acids Res 2008, 36 (22), 6959–6976. 10.1093/nar/gkn771.

(103) Isabelle, M.; Moreel, X.; Gagné, J.-P.; Rouleau, M.; Ethier, C.; Gagné, P.; Hendzel, M. J.; Poirier, G. G. Investigation of PARP-1, PARP-2, and PARG Interactomes by Affinity-Purification Mass Spectrometry. Proteome Sci 2010, 8, 22. 10.1186/1477-5956-8-22.

(104) Jungmichel, S.; Rosenthal, F.; Altmeyer, M.; Lukas, J.; Hottiger, M. O.; Nielsen, M. L. Proteome-Wide Identification of Poly(ADP-Ribosyl)Ation Targets in Different Genotoxic Stress Responses. Mol Cell 2013, 52 (2), 272–285. 10.1016/j.molcel.2013.08.026.

(105) Krietsch, J.; Rouleau, M.; Pic, É.; Ethier, C.; Dawson, T. M.; Dawson, V. L.; Masson, J.-Y.; Poirier, G. G.; Gagné, J.-P. Reprogramming Cellular Events by Poly(ADP-Ribose)-Binding Proteins. Mol Aspects Med 2013, 34 (6), 1066–1087. 10.1016/j.mam.2012.12.005.

(106) Mattsson, A.; Jernström, B.; Cotgreave, I. A.; Bajak, E. H2AX Phosphorylation in A549 Cells Induced by the Bulky and Stable DNA Adducts of Benzo[a]Pyrene and Dibenzo[a,l]Pyrene Diol Epoxides. Chem Biol Interact 2009, 177 (1), 40–47. 10.1016/j.cbi.2008.09.015.

(107) Lee, Y.-J.; Park, S.-J.; Ciccone, S. L. M.; Kim, C.-R.; Lee, S.-H. An in Vivo Analysis of MMC-Induced DNA Damage and Its Repair. Carcinogenesis 2006, 27 (3), 446–453. 10.1093/carcin/bgi254.

(108) Lemaire, M. A.; Schwartz, A.; Rahmouni, A. R.; Leng, M. Interstrand Cross-Links Are Preferentially Formed at the d(GC) Sites in the Reaction between Cis-Diamminedichloroplatinum (II) and DNA. Proc Natl Acad Sci U S A 1991, 88 (5), 1982–1985.

(109) Perry, R. P.; Kelley, D. E. Inhibition of RNA Synthesis by Actinomycin D: Characteristic Dose-Response of Different RNA Species. J Cell Physiol 1970, 76 (2), 127–139. 10.1002/jcp.1040760202.

(110) Jordan, P.; Carmo-Fonseca, M. Cisplatin Inhibits Synthesis of Ribosomal RNA in Vivo. Nucleic Acids Res 1998, 26 (12), 2831–2836.

(111) James, A.; Wang, Y.; Raje, H.; Rosby, R.; DiMario, P. Nucleolar Stress with and without P53. Nucleus 2014, 5 (5), 402–426. 10.4161/nucl.32235.

(112) Snodgrass, R. G.; Collier, A. C.; Coon, A. E.; Pritsos, C. A. Mitomycin C Inhibits Ribosomal RNA. J Biol Chem 2010, 285 (25), 19068–19075. 10.1074/jbc.M109.040477.

(113) Pontvianne, F.; Carpentier, M.-C.; Durut, N.; Pavlištová, V.; Jaške, K.; Schořová, Š.; Parrinello, H.; Rohmer, M.; Pikaard, C. S.; Fojtová, M.; Fajkus, J.; Saez-Vasquez, J. Identification of Nucleolus-Associated Chromatin Domains Reveals the Role of the Nucleolus in the 3D Organisation of the A. Thaliana Genome. Cell Rep 2016, 16 (6), 1574–1587. 10.1016/j.celrep.2016.07.016.

(114) Shav-Tal, Y.; Blechman, J.; Darzacq, X.; Montagna, C.; Dye, B. T.; Patton, J. G.; Singer, R. H.; Zipori, D. Dynamic Sorting of Nuclear Components into Distinct Nucleolar Caps during Transcriptional Inhibition. Mol Biol Cell 2005, 16 (5), 2395–2413. 10.1091/mbc.E04-11-0992.

(115) Walavalkar, K.; Gupta, S.; Kresoja-Rakic, J.; Raingeval, M.; Mungo, C.; Rubin, P.; Santoro, R. Single-Cell Dynamics of Genome-Nucleolus Interactions Captured by Nucleolar Laser Microdissection (NoLMseq). Nat Commun 2025, 16 (1), 11417. 10.1038/s41467-025-66294-7.

(116) Zhou, H.; Wang, Y.; Wang, Q.; Li, L.; Hu, Y.; Wu, Y.; Gautam, M.; Li, L. R-Loops Mediate Transcription-Associated Formation of Human rDNA Secondary Constrictions. J Cell Biochem 2021, 122 (10), 1517–1533. 10.1002/jcb.30074.

(117) Chan, Y. A.; Aristizabal, M. J.; Lu, P. Y. T.; Luo, Z.; Hamza, A.; Kobor, M. S.; Stirling, P. C.; Hieter, P. Genome-Wide Profiling of Yeast DNA:RNA Hybrid Prone Sites with DRIP-Chip. PLOS Genetics 2014, 10 (4), e1004288. 10.1371/journal.pgen.1004288.

(118) Sollier, J.; Cimprich, K. A. Breaking Bad: R-Loops and Genome Integrity. Trends Cell Biol 2015, 25 (9), 514–522. 10.1016/j.tcb.2015.05.003.

(119) Xiang, Y.; Laurent, B.; Hsu, C.-H.; Nachtergaele, S.; Lu, Z.; Sheng, W.; Xu, C.; Chen, H.; Ouyang, J.; Wang, S.; Ling, D.; Hsu, P.-H.; Zou, L.; Jambhekar, A.; He, C.; Shi, Y. RNA m6A Methylation Regulates the Ultraviolet-Induced DNA Damage Response. Nature 2017, 543 (7646), 573–576. 10.1038/nature21671.

(120) Tang, F.; Wang, Y.; Yuan, J.; He, X.; Zhang, Q.; Guo, S.; Dai, X.; Li, L.; Yuan, B.-F.; Wang, Y. Polymerase κ Recruits DDX23 To Promote R-Loop Resolution. Anal Chem 2026, 98 (15), 11564–11574. 10.1021/acs.analchem.6c01504.

(121) Simabuco, F. M.; Morello, L. G.; Aragão, A. Z. B.; Paes Leme, A. F.; Zanchin, N. I. T. Proteomic Characterization of the Human FTSJ3 Preribosomal Complexes. J Proteome Res 2012, 11 (6), 3112– 3126. 10.1021/pr201106n.

(122) Chen, H.; Yu, G.; Lin, H.; Zhou, X. A.; Liu, J.; Liu, H.; Luo, Y.; Cheng, K.; Mao, Z.; Ma, Y.; Wang, J.; Chen, K.; Wang, J.; Li, Y. Loss of FTSJ3 Promotes R-Loop-Associated DNA Damage and Facilitates Chemosensitivity in Lung Cancer Cells. Cancer Letters 2025, 631, 217877. 10.1016/j.canlet.2025.217877.

(123) Guo, C.; Tang, T.-S.; Bienko, M.; Dikic, I.; Friedberg, E. C. Requirements for the Interaction of Mouse Polkappa with Ubiquitin and Its Biological Significance. J Biol Chem 2008, 283 (8), 4658–4664. 10.1074/jbc.M709275200.

(124) Stenström, L.; Mahdessian, D.; Gnann, C.; Cesnik, A. J.; Ouyang, W.; Leonetti, M. D.; Uhlén, M.; Cuylen-Haering, S.; Thul, P. J.; Lundberg, E. Mapping the Nucleolar Proteome Reveals a Spatiotemporal Organization Related to Intrinsic Protein Disorder. Molecular Systems Biology 2020, 16 (8), e9469. 10.15252/msb.20209469.

